# PRC1 Sustains the Memory of Neuronal Fate Independent of PRC2 Function

**DOI:** 10.1101/2021.08.09.455667

**Authors:** Ayana Sawai, Sarah Pfennig, Milica Bulajić, Alexander Miller, Alireza Khodadadi-Jamayran, Esteban O. Mazzoni, Jeremy S. Dasen

**Affiliations:** Neuroscience Institute, Department of Neuroscience and Physiology, NYU School of Medicine, New York, NY 10016, USA; Department of Biology, New York University, New York, NY 10003, USA; Applied Bioinformatics Laboratories, Office of Science and Research, NYU School of Medicine, New York, NY 10016, USA

## Abstract

Polycomb repressive complexes (PRCs) 1 and 2 maintain stable cellular memories of early fate decisions by establishing heritable patterns of gene repression. PRCs repress transcription through histone modifications and chromatin compaction, but their roles in neuronal subtype diversification are poorly defined. We unexpectedly found that PRC2 is dispensable to preserve the morphogen-induced positional fates of spinal motor neurons (MNs), while PRC1 is essential for the specification of segmentally-restricted subtypes. Mutation of the core PRC1 component *Ring1* in mice leads to increased chromatin accessibility and ectopic expression of a broad variety of fates determinants, including Hox transcription factors, while neuronal class-specific features are maintained. Loss of MN subtype identities in *Ring1* mutants is due to the suppression of Hox networks by derepressed caudal *Hox* genes. These results indicate that PRC1 can function independently of *de novo* PRC2-dependent histone methylation to maintain chromatin topology and transcriptional memory at the time of neuronal differentiation.

## Introduction

Accurate control of gene expression is essential for the specification and maintenance of neural fates during development. Studies of cell type-restricted transcription factors have illuminated the mechanisms by which spatial and temporal regulation of gene expression gives rise to identifiable neuronal subtypes (Doe, 2017; Fishell and Kepecs, 2020; Hobert and Kratsios, 2019; Sagner and Briscoe, 2019; Venkatasubramanian and Mann, 2019). A parallel and critical mechanism of gene regulation is through the post-translational modification of histones, which enables and restricts transcription by modifying chromatin structure (Kishi and Gotoh, 2018; Schuettengruber et al., 2017). The Polycomb group (PcG) is key family of histone-associated proteins that play evolutionarily conserved roles in restricting gene expression during development (Blackledge et al., 2015; Gentile and Kmita, 2020; Simon and Kingston, 2009; Soshnikova and Duboule, 2009). In embryonic stem cells, cell fate determinants are repressed through PRC activities, and PRC-associated histone marks are subsequently removed from loci as cells differentiate (Boyer et al., 2006; Farcas et al., 2012; Tavares et al., 2012). PRC repression is maintained through cell division and after differentiation, and is thought to contribute to stable cellular memories of early patterning events (Ciabrelli et al., 2017; Coleman and Struhl, 2017). In vertebrates, much of our knowledge of how PRCs regulate gene expression has emerged from biochemical studies of PcG proteins or from the activity of these factors in proliferating cells. Despite an in depth understanding of the mechanisms of PRC action, how PcG proteins interact with gene regulatory networks in the CNS remains poorly understood.

The specification of neuronal fates in the vertebrate spinal cord provides a tractable system to elucidate the function of PcG proteins, as the pathways that determine identities are well-defined, and the molecular signatures of many subtypes are known (Butler and Bronner, 2015; Sagner and Briscoe, 2019). One neuronal class where fate specification has been closely examined is the spinal MN. A core set of transcription factors, including Hb9, Isl1/2, and Lhx3, determines class-specific features of MNs, including neurotransmitter identity (Shirasaki and Pfaff, 2002). The subsequent diversification of MNs into hundreds of muscle-specific subtypes is achieved through a conserved network of Hox transcription factors differentially expressed along the rostrocaudal axis (Philippidou and Dasen, 2013). During neural tube patterning, opposing gradients of retinoic acid (RA) and fibroblast growth factors (FGFs) provide spinal progenitors with a positional identity (Bel-Vialar et al., 2002; Dasen et al., 2003; Liu et al., 2001). These morphogens act, in part, by temporally and spatially depleting Polycomb-associated histone marks from *Hox* clusters (Mazzoni et al., 2013; Narendra et al., 2015). As progenitors exit the cell cycle, MNs continue to express *Hox* genes, where they regulate repertoires of subtype-specific genes (Catela et al., 2016; Dasen et al., 2005; Mendelsohn et al., 2017). Although the role of Hox proteins in the CNS are well-characterized (Parker and Krumlauf, 2020), and are known targets of PRC activities (Gentile and Kmita, 2020), the specific contributions of PRC1 and PRC2 to CNS maturation remain unclear, as few studies have directly compared their functions during embryonic development.

Polycomb repression is initiated by PRC2, which methylates histone H3 at lysine-27, permitting recruitment of PRC1 through subunits that recognize this mark, leading to chromatin compaction at genes targeted for repression (Margueron and Reinberg, 2011; Schuettengruber et al., 2017). PRC1 and PRC2 can also exist in a variety of configurations, which may contribute to neuronal subtype-specific activities. The core subunit of PRC1, Ring1, binds to one of six Polycomb group Ring finger (Pcgf) proteins (Gao et al., 2012). PRC1 containing Pcgf4 interacts with Cbx proteins (canonical PRC1) which recognize H3K27me3 (Bernstein et al., 2006; Morey et al., 2012). Variant forms of PRC1 containing Rybp can inhibit incorporation of Cbx proteins into PRC1, and bind target loci independent of H3K27me3 (Tavares et al., 2012; Wang et al., 2010). We previously found that a PRC1 component, Pcgf4, is required to establish rostral boundaries of *Hox* expression in differentiating MNs (Golden and Dasen, 2012). By contrast, deletion of a core PRC2 component, *Ezh2*, from MN progenitors had no apparent impact on fate specification, possibly due to compensation by *Ezh1* (Shen et al., 2008), or through use of variant PRC1 containing Rybp. These observations raise the question of what are the specific roles of canonical PRC1, variant PRC1, and PRC2 during MN subtype diversification.

To determine the function of PRCs during neural differentiation, we removed core components of these complexes from MN progenitors. We found that depletion of Ring1 proteins, essential constituents of all PRC1 isoforms, causes pronounced changes in transcription factor expression and a loss of Hox-dependent MN subtypes. Surprisingly, neither PRC2 nor variant PRC1 activities are required for rostrocaudal patterning at the time of MN differentiation. Deletion of *Ring1* leads to increased chromatin accessibility and derepression of a broad variety of cell fate determinants, while class-specific features of MNs are preserved. The derepression of caudal *Hox* genes in *Ring1* mutants leads to the suppression of MN subtype diversification programs. These findings indicate that PRC1 can preserve the memory of early patterning events in the CNS.

## Results

### PRC1 is Essential for Rostrocaudal Patterning During Neuronal Differentiation

To determine the relative contributions of PRC1 and PRC2 to neuronal specification, we analyzed mice in which core subunit-encoding genes were selectively removed from MN progenitors. *Ezh* genes encode the methyltransferase activity of PRC2, while *Eed* is required to enhance this function. We first generated mice in which both *Ezh* genes are conditionally deleted, by breeding *Ezh1* and *Ezh2* floxed lines to *Olig2::Cre* mice, which targets Cre to MN progenitors (*Ezh^MNΔ^* mice) (Hidalgo et al., 2012; Su et al., 2003). We confirmed MN-restricted loss of PRC2 activity by examining the pattern of H3K27me3 at E11.5, which was selectively depleted from progenitors and post-mitotic MNs (Figure 1A,B). Expression of Hb9 and *Vacht*, two general markers of MN identity, were grossly unchanged in *Ezh^MNΔ^* mice. (Figure 1C,D,G). We next analyzed expression of Hox proteins in *Ezh^MNΔ^* mice and found that the MN columnar subtype determinants Hoxc6, Hoxc9, and Hoxc10, were all expressed in their normal domains (Figure 1E,F, Supplementary Figure 1A-C). Similarly, we observed no changes in Hb9, Hoxc6, Hoxc9, and Hoxc10 expression in MNs after removal of the PRC2 component *Eed*, using *Sox1::Cre* (*Eed^NEΔ^* mice), which targets Cre to neuroectoderm (Supplementary Figure 1D-H) (Takashima et al., 2007; Yaghmaeian Salmani et al., 2018). Depletion of PRC2 activity from neural progenitors therefore does not affect MN class specification or rostrocaudal patterning.

**Figure 1.**
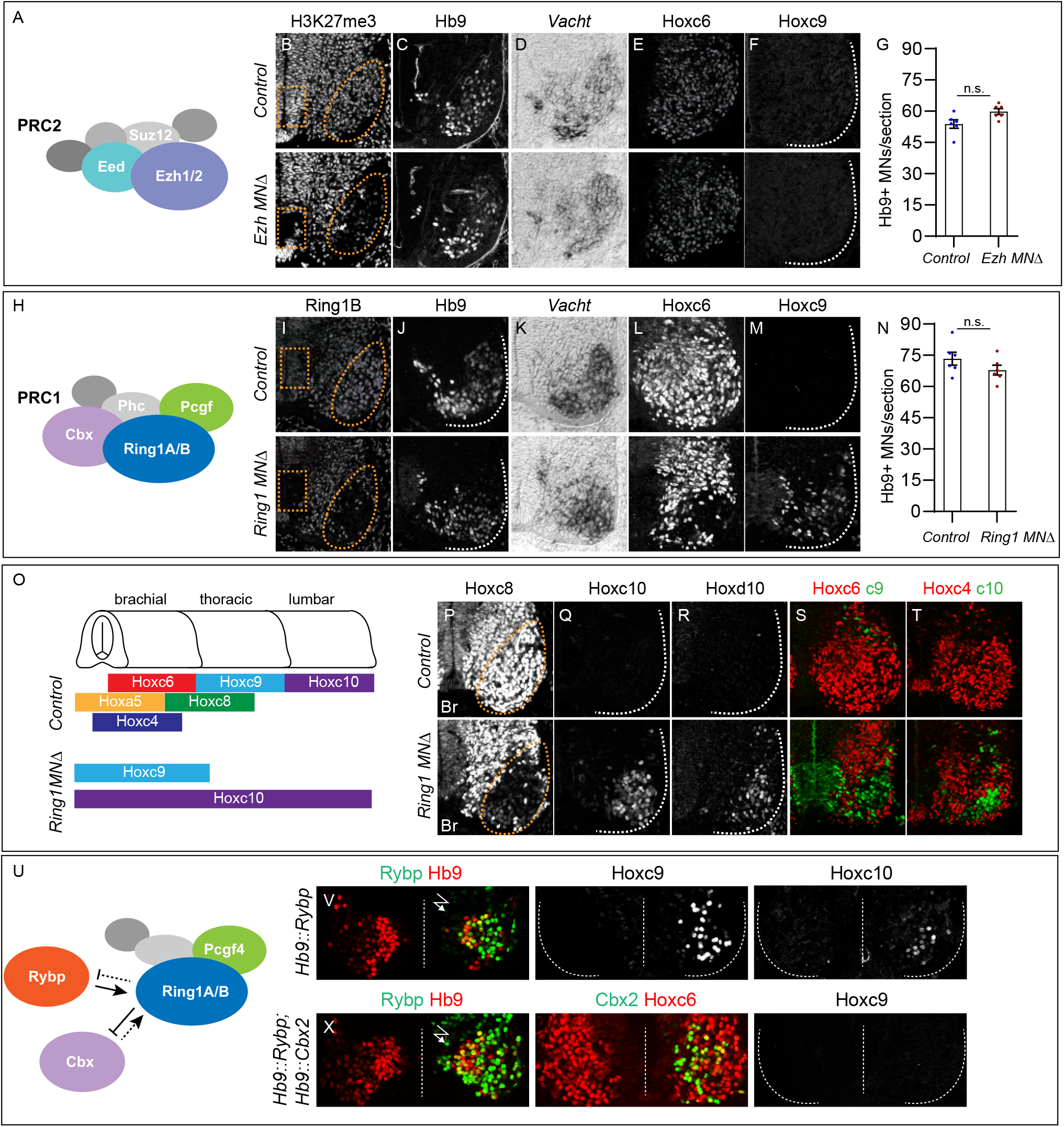
Roles of PRC1 and PRC2 in determining of *Hox* expression in spinal MNs. (A) Core components of PRC2. (B) Brachial spinal sections showing H3K27me3 is depleted from progenitors (boxed region) and post-mitotic MNs (oval) in E11.5 *EzhMN^Δ^* (*Ezh1^flox/flox^; Ezh2 ^flox/flox^, Olig2::Cre*^/+^) embryos. (C-D) MNs express Hb9 and *Vacht* in *Ezh^MNΔ^* mice. (E-F) Brachial Hoxc6 expression is normal in *Ezh^MNΔ^* mice, and no ectopic Hoxc9 is detected. (G) Quantification of MNs: 54+/−2 Hb9+ MNs per section in brachial controls, versus 60+/−1 in *Ezh^MNΔ^* mice, n=6 sections, p=0.0696. (H) Core components of PRC1. (I) Ring1B is selectively removed from progenitors (boxed region) and post-mitotic MNs (oval) in E12.5 *Ring1^MNΔ^* (*Ring1A^−/−^; Ring1B ^flox/flox^, Olig2::Cre*^/+^) mice. (J-K) *Ring1^MNΔ^* mice express Hb9 and *Vacht*. (L-M) Hoxc6 is lost from brachial MNs and Hoxc9 is ectopically expressed in *Ring1^MNΔ^* mice. (N) Quantification of MNs: 73+/-3 Hb9^+^ MNs in controls, versus 68+/−2 in *Ring1^MNΔ^* mice, n=6 sections, p=0.0956. (O) Summary of changes in Hox expression in MNs of *Ring1^MNΔ^* mice. (P-R) Loss of Hoxc8 and ectopic Hoxc10 and Hoxd10 expression in brachial MNs in *Ring1^MNΔ^* mice. (S-T) Colabeling of Hoxc6/Hoxc9 and Hoxc4/Hoxc10 in *Ring1^MNΔ^* mice, showing ectopically expressed caudal Hox proteins in place of rostral Hox protein expression in brachial segments. (U) Schematic of Cbx and Rybp interactions in PRC1. (V) Misexpression of Rybp in postmitotic MNs under *Hb9* in chick leads to ectopic Hoxc9 and Hoxc10 expression in brachial MNs. Bolt symbol indicates electroporated side of spinal cord. (X) Co-expression of Rybp and Cbx fails to induce Hoxc9 in brachial MNs. Panels B-F show brachial sections from E11.5 embryos; I-M, P-T brachial sections from E12.5 embryos; V,X brachial chick sections at HH st25. See also Supplementary Figure 1.

We next investigated the function of PRC1, which is thought to repress gene expression in a PRC2-dependent manner. We generated mice in which *Ring1B* is conditionally deleted from MN progenitors using *Olig2::Cre* in a global *Ring1A* mutant background (*Ring1^MNΔ^* mice) (Cales et al., 2008; del Mar Lorente et al., 2000). In *Ring1^MNΔ^* mice, Ring1B protein was selectively removed from progenitors and post-mitotic MNs (Figure 1H,I). The number of Hb9^+^ MNs and pattern of *Vacht* expression were similar to controls in *Ring1^MNΔ^* mice (Figure 1J,K,N). By contrast, Hox proteins normally expressed by forelimb-innervating brachial MNs were not detected, while thoracic and lumbar Hox proteins were ectopically expressed in more rostral MNs (Figure 1L,M,O-T, Supplementary Figure 1I,J,K). Brachial-level Hox proteins (Hoxc4, Hoxa5, Hoxc6 and Hoxc8), were selectively depleted from MNs, while thoracic and lumbar *Hox* determinants, (Hoxc9, Hoxc10, and Hoxd10) were derepressed in brachial segments (Figure 1Q-T). At thoracic levels Hoxc9 expression was attenuated and Hoxd10 was ectopically expressed (Supplementary Figure 1J). These results show that loss of *Ring1* leads to a marked rostral shift in *Hox* pattern, without affecting pan-MN molecular features (Figure 1O).

### Canonical PRC1 Regulates *Hox* Expression in MNs

Ring1 can interact with multiple Pcgf proteins, raising the question of which PRC1 configuration contributes to MN patterning. PRC1 containing Pcgf4 is required to establish *Hox* boundaries in MNs (Golden and Dasen, 2012), but can exist in two alternative configurations, depending on mutually exclusive incorporation of Cbx (canonical PRC1) or Rybp (variant PRC1) (Gao et al., 2012; Tavares et al., 2012). We examined the function of PRC1 isoforms first by manipulating Rybp and Cbx expression in MNs. We hypothesized that if canonical PRC1 regulates *Hox* expression, then overexpression of Rybp would inhibit binding of Cbx to Ring1, leading to MN phenotypes similar to *Ring1* mutants (Figure 1U). We used chick *in ovo* neural tube electroporation to express Rybp in postmitotic MNs using the *Hb9* promoter (*Hb9::mRybp*). Expression of *mRybp* under *Hb9* led to ectopic Hoxc9 and Hoxc10 expression at brachial levels (Figure 1V). If Rybp acts by displacing Cbx, then elevating Cbx levels should restore normal *Hox* expression. To test this, we co-electroporated *Hb9::mRybp* and *Hb9::mCbx2* at equivalent plasmid concentration, and no longer observed ectopic Hoxc9 in brachial segments (Figure 1X).

These observations are consistent with canonical PRC1 containing Pcgf4, Cbx, and Ring1 restricting *Hox* expression in MNs. Rybp is expressed by MNs at the time of their differentiation (Supplementary Figure 1L), raising the possibility that *Rybp* (or its paralog *Yaf2*) also plays regulatory roles during development. We therefore selectively deleted *Rybp* from MNs using *Olig2::Cre* mice in a *Yaf2*−/− background (*Rybp/Yaf2^MNΔ^* mice). Combined deletion of *Rybp* and *Yaf2* did not affect MN generation, Hoxc6, Hoxc9, or Hoxc10 expression (Supplementary Figure 1L). These findings indicate that canonical PRC1 maintains appropriate *Hox* expression, while variant PRC1 and PRC2 do not contribute to rostrocaudal patterning at the time of MN differentiation.

### PRC1 is Required for MN Subtype Diversification

Hox transcription factors play central roles in establishing neuronal subtype identities through regulating expression of subtype-specific genes. To determine the consequences of altered *Hox* expression in *Ring1^MNΔ^* mice, we analyzed the molecular profiles and peripheral innervation pattern of MNs. A key Hox target in MNs is the transcription factor *Foxp1*, which is essential for the differentiation of limb-innervating lateral motor column (LMC) and thoracic preganglionic column (PGC) neurons (Dasen et al., 2008; Rousso et al., 2008). In *Ring1^MNΔ^* mice, Foxp1 expression is lost from brachial and thoracic MNs, and markedly reduced in lumbar MNs (Figure 2A,B,G, Supplementary Figure 2C). By contrast, in *Ezh^MNΔ^*, *Eed^NEΔ^* and *Rybp/Yaf2^MNΔ^* mice, Foxp1 expression was maintained, consistent with the preservation of normal *Hox* profiles in these mutants (Supplementary Figure 1L, 2A,B). In *Ring1^MNΔ^* mice, expression of the LMC marker Raldh2 was lost at brachial levels, and the thoracic PGC marker nNos was not detected (Figure 2C, D). Moreover, expression of determinants of respiratory phrenic MNs (Scip^+^ Isl1/2^+^), was markedly depleted, and likely contributes to the perinatal lethality of *Ring1^MNΔ^* mice (Supplementary Figure 2D). These results indicate that *Ring1* deletion leads to a loss of genes acting downstream of Hox function in MNs.

**Figure 2.**
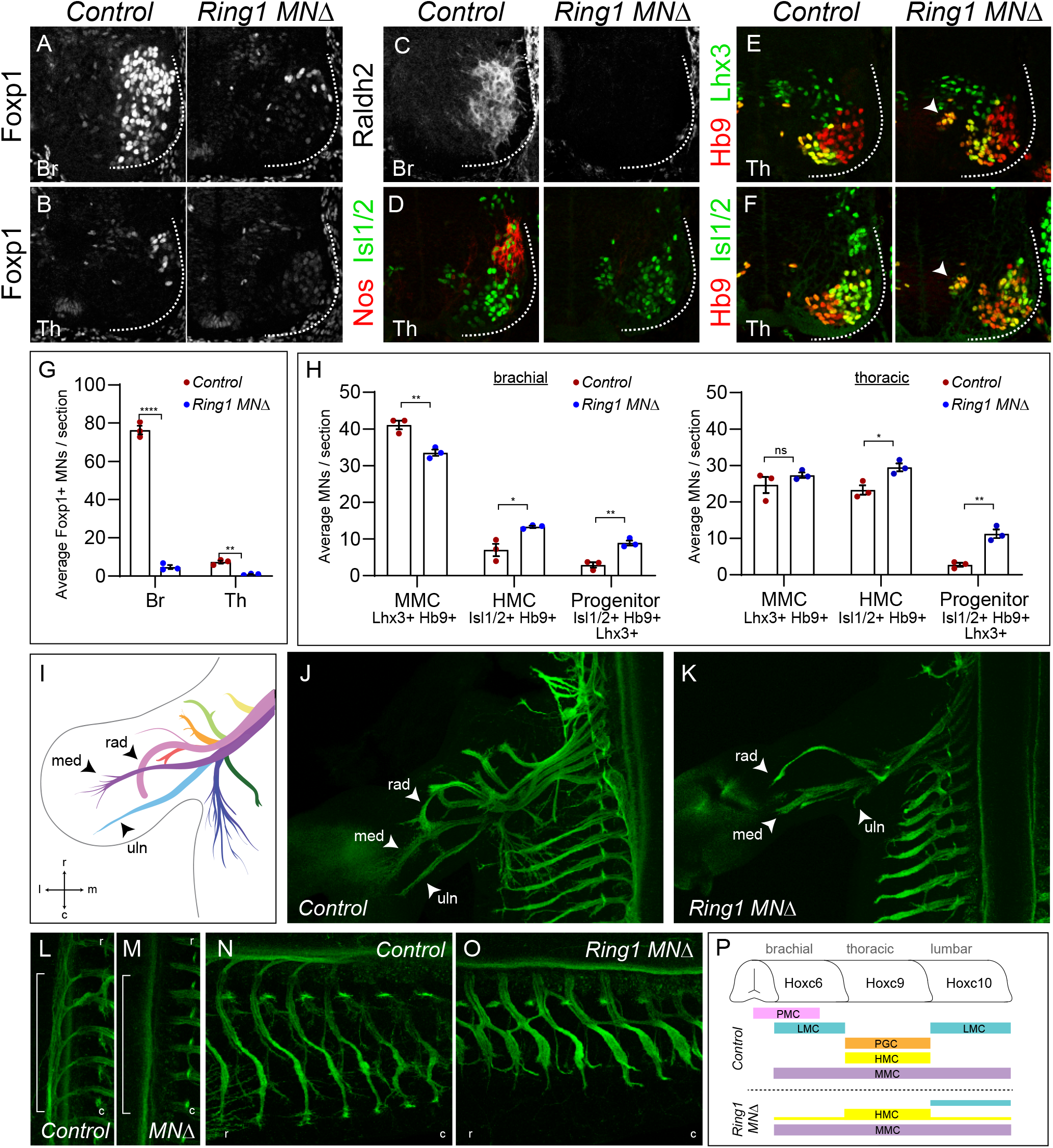
*Ring1* is essential for the specification of Hox-dependent MN subtypes. (A-B) Foxp1 expression is reduced in brachial (Br) and thoracic (Th) segments of *Ring1^MNΔ^* mice at E12.5. (C-D) Expression of the brachial LMC marker Raldh2 and thoracic PGC marker nNos were lost in *Ring1^MNΔ^* mice. (E-F) Staining of Hb9^+^, Lhx3^+^ (MMC) and Hb9, Isl1/2^+^ (HMC) neurons. In *Ring1^MNΔ^* mice, we also observed a population of medial neurons that coexpressed Isl1/2, Hb9, and Lhx3 (indicated by arrow heads). (G) Quantification of Foxp1 reduction. (H) Quantification of MMC (Hb9^+^, Lhx3^+^), HMC (Isl1/2^+^, Hb9^+^, Lhx3^+^) and medial “progenitors” (Isl1/2^+^, Hb9^+^, Lhx3^+^) MNs (arrow heads in E,G. Panels G-H show average from n=3 mice, 4 sections each animal. Data shown in graphs shown as mean± SEM. *p<0.05, **p<0.01, ****p<0.0001. (I) Schematic of nine primary nerves in E12.5 mouse forelimb (Adapted from Catela et al., 2016) med=median, rad=radial, uln=ulnar nerves. Rostral (r), caudal (c), medial (m), and lateral (l) orientation shown. (J-K) Forelimb motor axons of an E12.5 control and *Ring1^MNΔ^* mouse labeled by *Hb9::GFP*. (L-M) Innervation of sympathetic chain ganglia (from PGC neurons) in control and *Ring1^MNΔ^* mice. Bracket shows region of PGC projections along rostrocaudal axis. (N-O) Innervation of dorsal and ventral axial muscles by MMC and HMC respectively. In *Ring1^MNΔ^* mice, axial motor projections are shorter and thicker. (P) Summary of MN columnar organization of control and *Ring1^MNΔ^* mice. See also Supplementary Figure 2.

We next assessed the impact of *Ring1* deletion on two MN subtypes that are specified independent of Hox function, the hypaxial and median motor columns (HMC and MMC). In *Ring1^MNΔ^* mice, MMC neurons (Lhx3^+^ Hb9^+^) were generated at normal numbers in thoracic segments, while the number of HMC neurons (Isl1/2^+^ Hb9^+^ Lhx3^−^) was increased in thoracic and brachial segments (Figure 2E,F,H, Supplementary Figure 2E,F). The increase in HMC neurons in *Ring1* mutants is likely due to a reversion of presumptive Hox-dependent subtypes to an HMC fate, similar to mice in which Hox function is disrupted (Dasen et al., 2008; Hanley et al., 2016). Mutation in *Ring1* therefore depletes Hox-dependent subtypes, with the remaining MNs having a more ancestral axial identity (Figure 2P).

As we observed a dramatic loss of segmentally-restricted MN subtypes in *Ring1^MNΔ^* mice, we next assessed the impact on peripheral innervation pattern. To trace motor axon projections, we crossed *Ring1^MNΔ^* mice to an *Hb9::GFP* reporter, in which all MN axons are labelled with GFP. In control mice there are nine primary trajectories of forelimb-innervating motor axon at E12.5 (Figure 2I,J) (Catela et al., 2016). In *Ring1^MNΔ^* mice, only three nerve branches, radial, median, and ulnar were visible but appeared prematurely truncated and unbranched (Figure 2K). In the trunk, innervation of sympathetic chain ganglia was lost, consistent with a loss of PGC fates (Figure 2L,M). Projections to dorsal and ventral axial muscles by MMC and HMC subtypes were maintained in *Ring1^MNΔ^* mice, although axons were shorter than in controls (Figure 2N, O). Loss of *Ring1* therefore causes severe defects in innervation pattern, while the trajectories of axial MNs are relatively spared.

### Loss of *Ring1* Causes Broad Derepression of Developmental Fate Determinants

Polycomb proteins regulate diverse aspects of differentiation by restricting gene expression during development. To investigate changes in gene expression after deletion of *Ring1* genes in an unbiased manner, we performed RNAseq on MNs isolated from *Ring1^MNΔ^* mice. We purified MNs from control and *Ring1^MNΔ^*;*Hb9::GFP* embryos at E12.5 by flow cytometry (n=4 *Ring1* mutants, and n=4 Cre^−^ controls), and performed RNAseq. Because *Ring1* mutants display segment-specific phenotypes, we collected MNs from brachial, thoracic, and lumbar levels and profiled each population independently.

We identified a total of 1001 upregulated and 641 downregulated genes in *Ring1^MNΔ^* mice (log_2_-FC>2, FDR<0.1), with 391 genes upregulated in all three segmental regions, compared to 46 genes that were commonly downregulated (Figure 3A-C, Supplementary Figure 3A,C, Table 1). Thus, many upregulated genes are shared between each segment, while downregulated genes tended to be segment-specific. Strikingly, 76% of the top 100 derepressed genes (by log_2_-FC) at brachial levels encode transcription factors, and include genes normally involved in the specification of spinal interneurons (e.g. *Gata3*, *Shox2*, *Lhx2*), brain regions (*FoxG1*, *Six6, Lhx8*), and non-neuronal lineages (Figure 3B).

**Figure 3.**
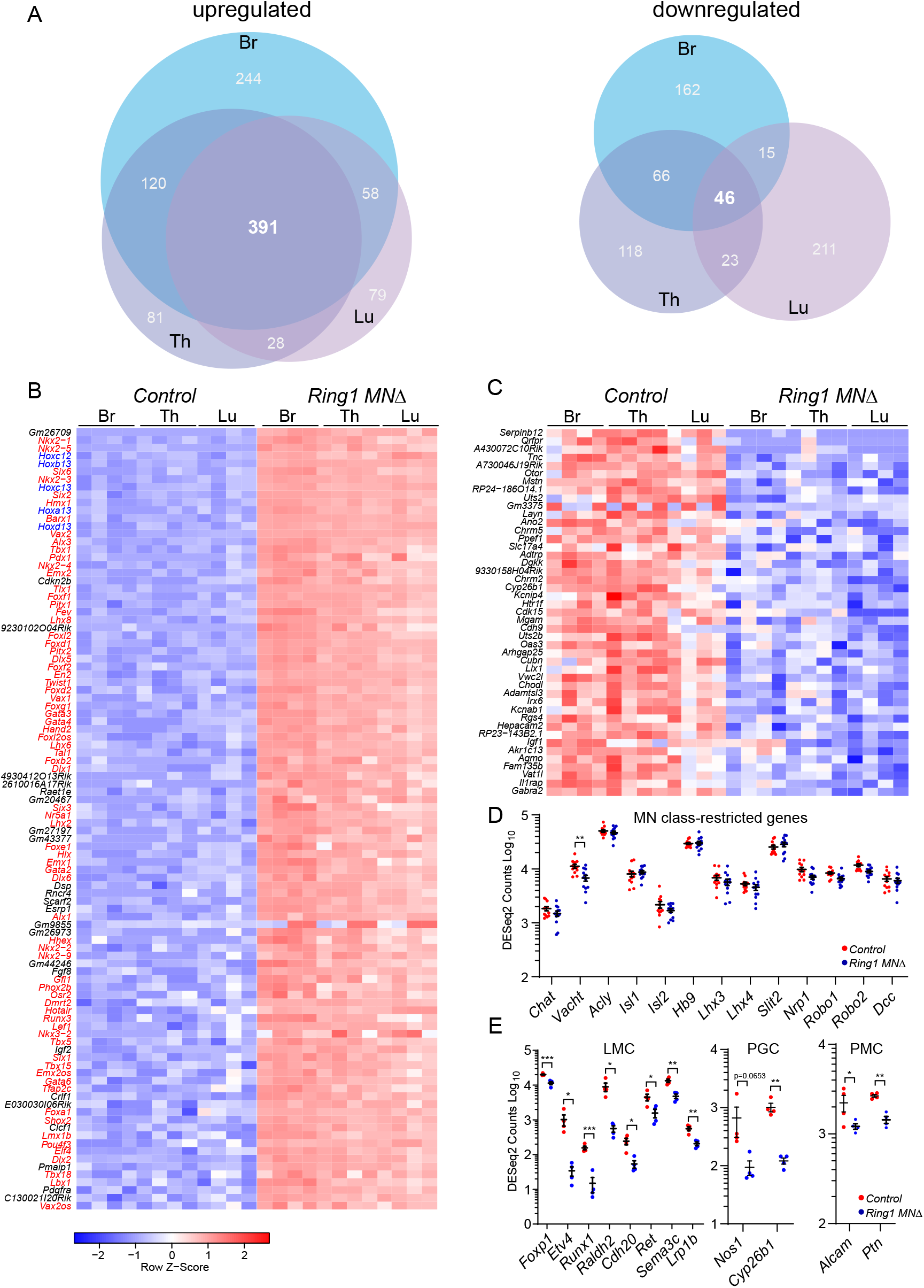
*Ring1* is required to restrict transcription factor expression in spinal MNs. (A) Venn diagrams showing the number of upregulated (left) and downregulated (right) genes in bracial (Br), thoracic (Th), and lumbar (Lu) MNs upon loss of *Ring1* from MNs by RNAseq (log_2_-FC>2, FDR<0.1). (B) Heat map of top 100 upregulated genes (by log_2_-FC) in *Ring1^MNΔ^* mice and control MNs. Genes shown in blue are *Hox* genes and other transcription factor are shown in red. (C) Heat map of 45 genes downregulated in *Ring1^MNΔ^* mice in comparison to control MNs. (D) Plots of DESeq2 counts of genes associated with MN class identity. Each data point shows DESeq2 counts for each sample, and segment-specific counts are plotted together. Expression of *Slc18a3 (Vacht)* is reduced (padj.=0.026477) in *Ring1^MNΔ^*. (E) DESeq2 counts of genes associated with specific MN subtype identities were reduced in *Ring1^MNΔ^* mice. Counts for LMC and PMC markers are from Br segments, PGC from Th segments. Black bars shown in graphs indicate mean± SEM. *p<0.05, **p<0.01, ***p<0.001, ****p<0.0001. See also Supplementary Figure 3.

Although a variety of cell fate determinants were derepressed in *Ring1^MNΔ^* mice, our RNAseq analyses provide further evidence that core molecular determinants of MN class identity are preserved in *Ring1* mutants. We found no significant changes in transcription factors (*Isl1*, *Isl2*, *Hb9*, *Lhx3*, *Lhx4*), guidance molecules (*Slit2*, *Robo1*, *Robo2*, *DCC*) and neurotransmitter genes (*Chat*, *Acly*) associated with pan-MN features (Figure 3D). There was a modest decrease in expression *Slc18a3 (Vacht)* (Figure 3D). Expression of genes that mark other excitatory (*Slc17a7*, *Slc17a6*) or inhibitory (*Slc32a1, Gad2*, *Gad1*,) neuronal classes were not markedly changed in *Ring1* mutants (Supplementary Figure 3B). By contrast, expression of genes associated with MN subtype identities was decreased in *Ring1^MNΔ^* mice, including genes that mark limb-innervating (e.g *Foxp1*, *Etv4, Raldh2, Runx1*), thoracic-specific (*Nos1*, *Cyp26b*), and respiratory (*Alcam*, *Ptn*) subtypes (Figure 3E).

To validate these changes in gene expression, we performed mRNA *in situ* hybridization on a subset of upregulated or downregulated genes. We found that *Gata3*, *Lhx8*, and *Lhx2* were markedly upregulated in MNs of *Ring1^MNΔ^* mice (Figure 4A-F), while other genes (*Foxg1*, *Six6*, *Pitx2, Shox2, Nkx2.1*) showed less prominent, but detectable, ectopic expression (Supplementary Figure 4A-J). We also examined expression of previously uncharacterized genes that were downregulated in all three segmental regions. We found *Rgs4*, *Uts2b*, *Vat1*, and *Gabra2* were expressed by MNs of control embryos, and were downregulated in *Ring1* mutants (Figure 4G-L, Supplementary Figure 4K,L). These results indicate that despite the derepression of multiple fate determinants in *Ring1^MNΔ^* mice, core features of MN identity are preserved, but that subtype diversification programs are selectively disrupted.

**Figure 4.**
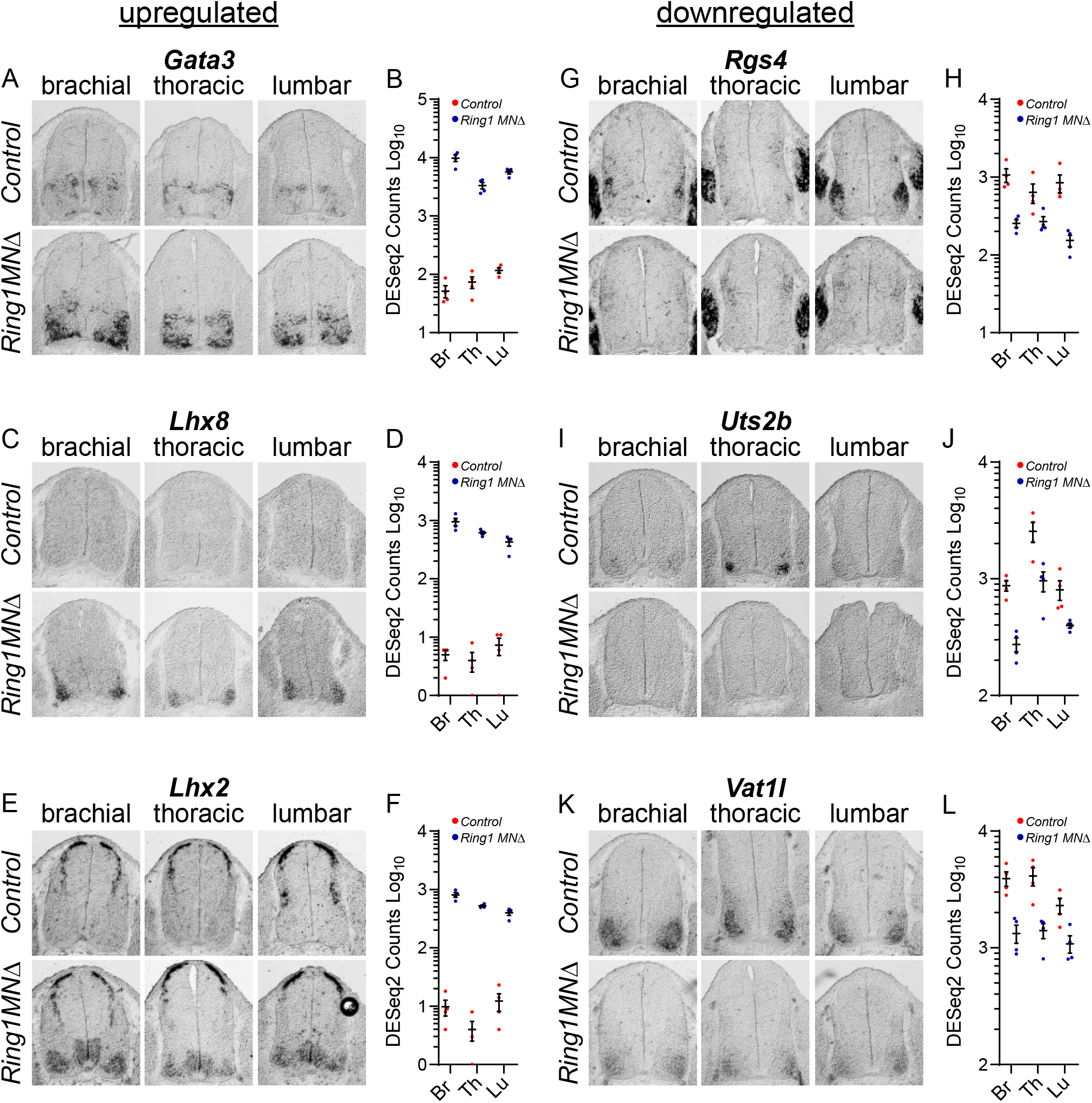
Analyses of misregulated genes in *Ring1* mutants. (A,C,E) *In situ* mRNA hybridization of selected upregulated genes from *Ring1^MNΔ^* RNAseq. Images show sections of brachial, thoracic, and lumbar segments from E12.5 control and *Ring1^MNΔ^* mice. (B,D,F) Graphs of DESeq2 counts for each upregulated gene in each segment. Data points show DESeq2 counts from MNs of individual animals from indicated segments. (G, I, K) Analysis of downregulated genes by *in situ* hybridization. *Rgs4* and *Uts2b* displayed elevated expression in specific segmental levels in controls, suggesting that a subset of the commonly-downregulated genes are also Hox-dependent. *Rgs4* expression is normally elevated in LMC neurons (panel G), while *Uts2b* is elevated in thoracic segments of controls (panel I) (H,J,L) Graphs of DESeq2 counts for each downregulated gene in each segment. See also Supplementary Figure 4.

### Selective Derepression of Caudal *Hox* genes in *Ring1^MNΔ^* mice

As our preliminary analyses revealed altered expression in a subset of *Hox* genes in *Ring1^MNΔ^* mice, we further evaluated *Hox* genes in our RNAseq dataset. Consistent with the analysis of Hox protein expression, rostral *Hox* genes were reduced in brachial segments of *Ring1^MNΔ^* mice, while *Hox10-Hox13* genes were derepressed (Figure 5A, Supplementary Figure 5A-C). We observed derepression of caudal *Hox* genes from each of the four vertebrate *Hox* clusters, including genes not normally detectable in the ventral spinal cord (e.g *HoxB* genes) (Dasen et al., 2005) (Figure 5A, Supplementary Figure 5A-C). The extent of caudal *Hox* derepression correlated with the relative position of genes within a cluster, with *Hox13* genes displaying the most pronounced derepression in *Ring1^MNΔ^* mice (by FC), relative to other caudal *Hox* genes (Figure 5B, Supplementary Figure 3C). The marked derepression of *Hox13* genes was observed in each of the three segmental levels we analyzed, while *Hox10* genes were selectively derepressed in brachial and thoracic segments (Figure 5A, Supplementary Figure 5A-C).

**Figure 5.**
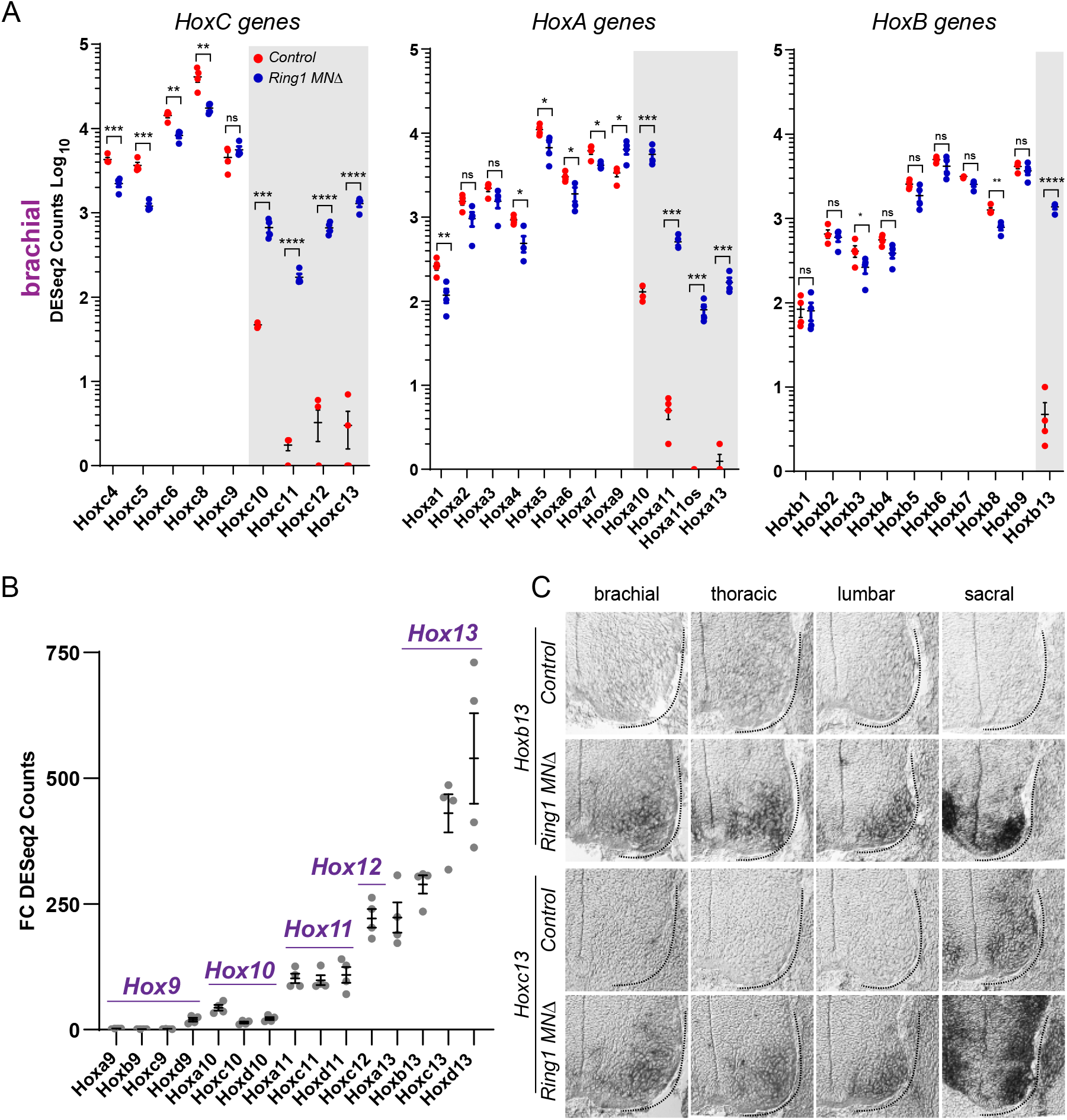
Derepression of caudal *Hox* genes in *Ring1* mutants. (A) DESeq2 counts of *HoxC*, *HoxA,* and *HoxB* cluster genes in brachial segments in control and *Ring1^MNΔ^* mice showing derepression of caudal *Hox* genes. Grey shaded regions highlight *Hox* genes that are derepressed in *Ring1^MNΔ^* mice. *Hoxc9* does not show significant derepression, likely because it is normally expressed by caudal brachial MNs. Black bars shown in graphs indicate mean± SEM. *p<0.05, **p<0.01, ***p<0.001, ****p<0.0001. (B) Comparison of *Hox9-Hox13* paralog gene derepression in brachial segments. Graph shows absolute fold changes of DESeq2 counts, showing a marked increase for caudal *Hox13* paralogs in *Ring1^MNΔ^* mice. Each data point shows individual counts for *Ring1* mutants/average of controls. (C) *In situ* of *Hoxb13* and *Hoxc13* mRNA transcripts in E12.5 embryos. *Hoxb13* is normally not detectable in spinal cord, but is derepressed in MNs in *Ring1^MNΔ^* mice. *Hoxc13* transcripts are normally restricted to sacral segments but derepressed in rostral segments in *Ring1^MNΔ^* mice. See also Supplementary Figure 5.

To further validate these findings, we analyzed *Hox13* expression by *in situ* hybridization. In controls *Hoxc13* is restricted to sacral segments, while *Hoxb13* is not detected in MNs (Figure 5C). In *Ring1^MNΔ^* mice, both *Hoxb13* and *Hoxc13*, were de-repressed in MNs throughout the rostrocaudal axis (Figure 5C). By contrast, *in situ* hybridization of *Hoxc6* and *Hoxc9* expression revealed reduced expression at brachial and thoracic levels, respectively (Supplementary Figure 5C). Thus, ectopic expression of *Hox13* genes in *Ring1* mutants is associated with a loss of rostral *Hox* gene expression.

### *Ring1* is Essential to Maintain MN Chromatin Topology

Our findings indicate that in the absence of *Ring1* genes, a broad variety of cell fate determinants are ectopically expressed in MNs, while only a subset of caudal *Hox* genes are derepressed. As PRC1 restricts gene expression through chromatin compaction, we investigated whether removal of *Ring1* leads to changes in DNA accessibility at derepressed loci. We used Assay for Transposase-Accessible Chromatin with high-throughput sequencing (ATACseq) to identify genomic regions which have gained or lost accessibility in *Ring1^MNΔ^* mice. We purified *Hb9::GFP^+^* MNs at E12.5 from control and *Ring1^MNΔ^* embryos at brachial, thoracic, and lumbar levels, and performed ATACseq. In control samples, we observed a progressive opening of caudal *Hox* genes in more rostral segments. For example, the accessibility of *Hox9* genes increases from brachial to thoracic segments, while *Hox10* and *Hox11* genes are more accessible in lumbar segments (Supplementary Figure 6A, B).

To determine which genomic regions gained accessibility in *Ring1* mutants, we compared ATACseq profiles between MNs of control and *Ring1^MNΔ^* mice. We identified a total of 2,305 loci that gained accessibility in *Ring1^MNΔ^* embryos, 324 (14%) of which were common to all three segmental levels (Figure 6A, Table 2). Common genes included caudal *Hox* genes (*Hoxa13*, *Hoxc13*) and other transcription factors (e.g. *Foxg1*, *Lhx2, Pitx2*) that were derepressed in our RNAseq analyses (Figure 6C). We also identified 1,264 loci that lost accessibility in *Ring1^MNΔ^* mice, 39 (3%) of which were common to all three segments (Figure 6A). Thus, similar to our RNAseq results, loss of *Ring1* leads to increased accessibility in many genes that are shared among each segment, while genes that lose accessibility tend to be segment-specific.

To assess which genes may be directly repressed by PRC1, we examined the overlap between transcripts that were upregulated and loci that gained accessibility in *Ring1^MNΔ^* mice. We found that 18% (148/813) of genes that were ectopically expressed at brachial segments also gained accessibility in *Ring1^MNΔ^* mice (Figure 6B). We found 19 genes, including *Hoxa13* and *Hoxc13*, were derepressed and gained accessibly in all three segments (Supplementary Figure 6C). Since lumbar segments still retain features of Hox-dependent subtypes, we also compared the overlap between genes that were upregulated and gained accessibility in brachial and thoracic MNs. We identified 73 genes, 56 (77%) of which encode transcription factors, including each of the caudal *Hox* genes we found by RNAseq (Figure 6B, Supplementary Figure 6C). The gain of accessibility at transcription factor-encoding genes was prominent near transcription start sites (Figure 6C), and regions that gained accessibility in *Ring1^MNΔ^* neurons correspond to regions shown to be bound by Ring1B in ES cells (Supplementary Figure 6D) (Bonev et al., 2017).

In agreement with our RNAseq analysis, *Hox13* paralogs showed pronounced increases in accessibility, with each of the four *Hox13* genes gaining accessibility in each of the three segmental levels (Figure 6D,E). By contrast, rostral *Hox4*-*Hox8* genes, which are transcriptionally downregulated in *Ring1* mutants, were not among that targets that lost accessibility (Figure 6D). These results suggest that reduced expression of rostral *Hox4-Hox9* genes in *Ring1* mutants is not due to a loss in chromatin accessibility.

### Hox13 Paralogs Represses Rostral *Hox* genes by Engaging Accessible Chromatin Domains

As PRC1-mediated chromatin compaction is essential for *Hox* repression, it is surprising that *Hox4-Hox9* genes are diminished in *Ring1* mutants, without a noticeable reduction in chromatin accessibility at these loci (Supplementary Figure 6D). Because cross-repressive interactions between *Hox* genes themselves are an important regulatory mechanism determining *Hox* boundaries (Philippidou and Dasen, 2013), it is possible that ectopically expressed *Hox13* genes directly repress multiple *Hox* genes in *Ring1^MNΔ^* mice. To test this, we first analyzed MNs derived from ES cells (ESC-MNs) in which *Hoxc13* expression can be induced upon doxycycline (Dox) treatment (Bulajic et al., 2020). ESC-MNs differentiated via RA and Sonic Hedgehog agonist are characterized by expression of *Hox4-Hox5* paralogs. RNAseq analyses revealed that *Hoxa4*, *Hoxa5*, *Hoxc4*, and *Hoxc5* expression were markedly reduced after Dox-induced *Hoxc13* expression in MN progenitors, compared to non-induced ESC-MNs (Figure 7A, Supplementary Figure 7A-C). This repressive effect also appears to be direct, as ChIPseq analysis indicates that Hoxc13 can bind at multiple *Hox* genes (Figure 7A).

**Figure 7.**
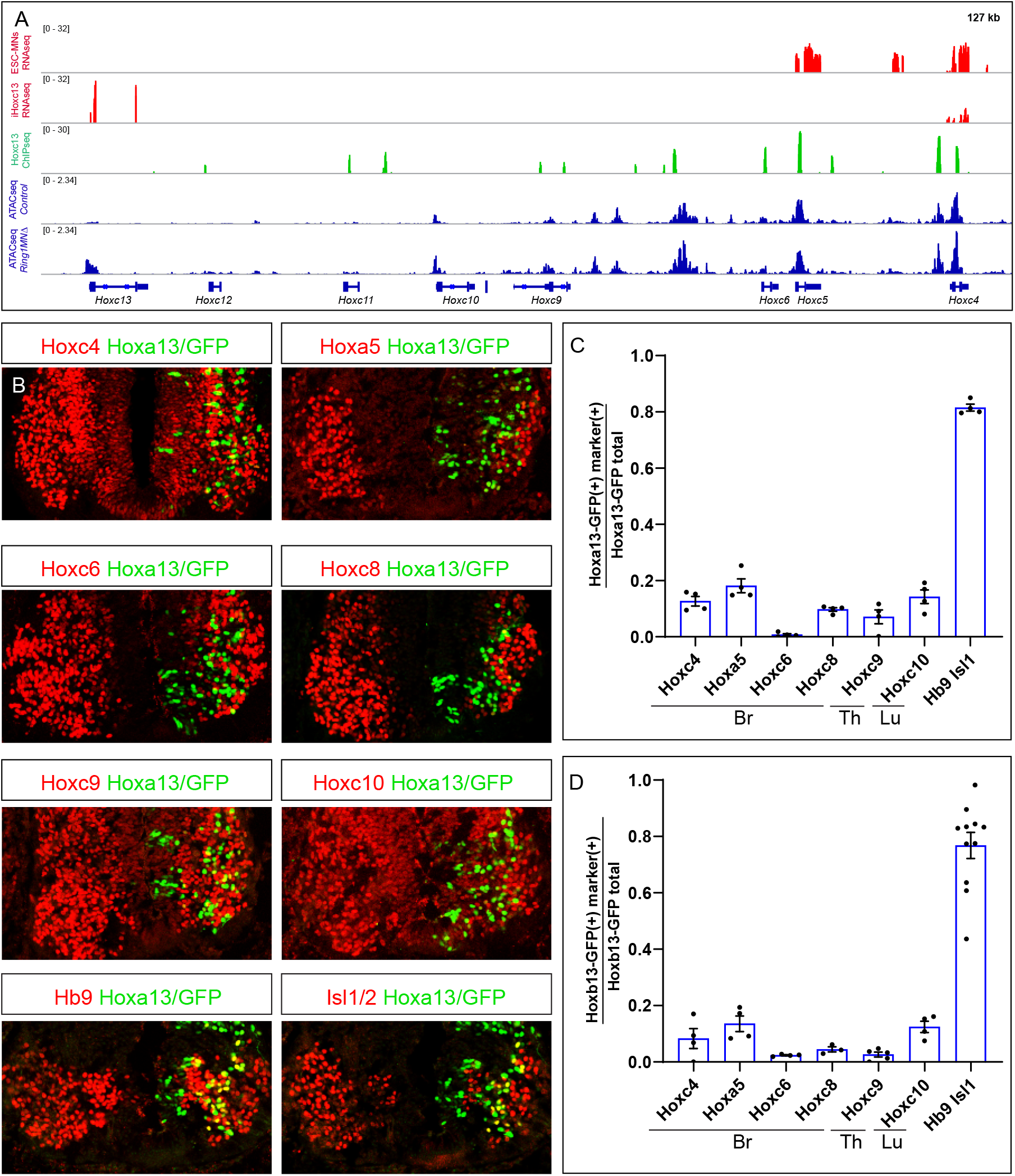
Hox13 proteins repress multiple *Hox4-Hox10* paralogs. (A) Effects of *Hoxc13* induction in ESC-MNs. Top panels show IGV browser views (log scale) of RNAseq (red) and ChIPseq tracks (green) at the *HoxC* cluster. Induced Hoxc13 binds near multiple *HoxC* genes and represses expression of *Hoxc4* and *Hoxc5*. Bottom panels show ATACseq in brachial control and *Ring1* mutant MNs at the *HoxC* cluster (blue). (B) Misexpression of *mHoxa13-ires-nucGFP* represses more rostral *Hox* genes. Hoxc4, Hoxa5, Hoxc6 and Hoxc8 were analyzed in brachial segments, Hoxc9 in thoracic segments, and Hoxc10 in lumbar segments. Hb9 and Isl1 images are from brachial segments. (C,D) Quantification of percentage of electroporated cells which retained the expression of indicated Hox protein and MN markers upon *Hoxa13* or *Hoxb13* misexpression. Quantified electroporated cells were selected from the ventrolateral spinal cord, where MNs reside. Hb9 and Isl1/2 quantification shows percentage of GFP^+^ neurons that express either marker, and are shown for Br segments in panel B, and Br and Th segments in C. Data in graphs shown as mean± SEM, averaged from n=3 animals, 4 sections each. See also Supplementary Figure 7.

Hox13 proteins can target inaccessible chromatin (Bulajic et al., 2020; Desanlis et al., 2020), while repression by the *Drosophila* Hox protein Ubx is associated with chromatin compaction (Loker et al., 2021). To examine the possible effects of Hox13 proteins on local chromatin structure, we compared accessibility at *Hox* loci in both control and *Ring1* mutants, relative to the location of Hoxc13 binding sites. At *Hox4-Hox5* genes, Hoxc13 sites correspond to regions that are accessible in both control and *Ring1* mutant MNs (Figure 7A, Supplementary Figure 7A-C). Hox13 proteins therefore appear to repress rostral *Hox* genes, in part, through binding at pre-existing accessible regions.

To test whether *Hox13* genes can repress multiple *Hox* paralogs *in vivo*, we used chick neural tube electroporation to express *Hox13* genes within brachial, thoracic, and lumbar segments, and assessed *Hox4-Hox10* gene expression. We electroporated *pCAGGs::mHoxa13-ires-nucGFP* or *pCAGGs::mHoxb13-ires-nucGFP* and found that both cell-autonomously repressed expression of brachial, thoracic, and lumbar Hox proteins (Hoxc4, Hoxa5, Hoxc6, Hoxc8, Hoxc9, and Hoxc10) (Figure 7B-D). By contrast, misexpression of *Hox13* genes did not affect expression of the general MN markers Isl1/2 and Hb9 (Figure 7B-D). These results indicate that Hox13 proteins can directly repress multiple *Hox* genes, and likely contribute to the MN fate specification defects of *Ring1^MNΔ^* mice.

## Discussion

The Polycomb group encompass a large and diverse family of proteins essential for maintaining epigenetic memory of early patterning events. Classically, PcG-mediated repression is thought to depend on recruitment of PRC1 through recognition of histone methylation marks deposited by PRC2 activity. Although alternative, H3K27me3-independent, mechanisms of PRC1 repression have been described (Gao et al., 2012; Tavares et al., 2012), the relative contribution of PRC1 and PRC2 to neuronal fate specification have not been resolved. In this study, we found that genetic depletion of PRC2 components has no discernable impact on neural class or subtype differentiation programs at the time of MN differentiation. By contrast, a core PRC1 subunit, Ring1, is required to restrict transcription factor expression in the CNS, maintain neuronal subtype-specific chromatin topology, and determine rostrocaudal boundaries of *Hox* expression. Our findings indicate PRC1 plays an key role in regulating expression of cell fate determinants during MN differentiation, safeguarding neurons from acquiring inappropriate gene regulatory programs.

### PRC Functions in Neural Development

Polycomb proteins function in diverse aspects of CNS maturation, including the temporal transition from neurogenesis to gliogenesis, maintenance of adult neural stem cell fates, and restriction of gene expression in mature neurons (Di Meglio et al., 2013; Fasano et al., 2009; Hirabayashi et al., 2009; Molofsky et al., 2003; von Schimmelmann et al., 2016). The role of PRCs in neural subtype diversification has been challenging to study, in part, due the complexity and dynamic temporal regulation of its constituents. By selectively removing core PRC1 and PRC2 subunits from neural progenitors, we determined the relative contributions of these chromatin-associated complexes to neuronal diversification. We found that PRC1 represses a broad variety of cell fate determinants, with the majority of highly derepressed genes in *Ring1* mutants encoding transcription factors. Ectopic expression of Ring1-regulated targets is associated with a disruption in chromatin topology, resulting in a gain in accessibility of derepressed genes.

Despite the derepression of multiple fate determinants in the absence of PRC1 function, expression of transcription factors that define MN class identity (e.g. Hb9, Isl1/2, and Lhx3), are largely unchanged, and MNs do not acquire characteristics of other neuronal classes. This observation suggests that once a class identity has been established, ectopic expression of fate determinants is insufficient to respecify basic neuronal features, such as neurotransmitter identity. In *C. elegans* the ability of transcription factors to reprogram neuronal class identity requires removal of histone deacetylase proteins (Tursun et al., 2011), suggesting that additional regulatory constraints, such as histone acetylation and/or DNA-methylation, restricts the ability of ectopic transcription factors to interfere with class-specific programs in *Ring1* mutants.

While basic elements of MN fate are maintained in the absence of PRC1, their differentiation into molecularly distinct subtypes is severely compromised. The loss of subtype-specific programs in brachial and thoracic segments can be attributed to the attenuation of *Hox4-Hox9* paralog expression, as mice lacking these *Hox* genes show a similar loss in MN subtype features (Jung et al., 2010; Jung et al., 2014; Philippidou et al., 2012). Selective loss of *Hox* function in MNs also appears to account for the observation that the majority of downregulated genes in *Ring1* mutants are segment-specific.

### PRC-Dependent and Independent Mechanisms of Rostrocaudal Patterning in the CNS

We found that removal of PRC1 leads to the derepression and increased accessibility of caudal *Hox* genes. Loss of PRC1 does not result in a sustained derepression of all *Hox* genes, as *Hox* genes normally expressed by brachial- and thoracic-level MNs are attenuated in the absence of *Ring1*. These findings are reminiscent of the function of PRCs in *Drosophila*, where loss of PRC function leads to ectopic expression of the caudal *Hox* gene *Abd-b* and repression of *Ubx* expression (Struhl and Akam, 1985; Struhl and White, 1985), likely reflecting cross-repressive interactions between caudal Hox proteins and rostral *Hox* genes. Notably, reduced rostral *Hox* gene expression in *Ring1* mutants is not a reflection of reduced chromatin accessibility at these targets, consistent with a model in which this form of repression is not associated with global changes in chromatin structure.

In *Ring1* mutants, *Hox13* genes are derepressed in MNs, and appear to contribute to the loss of MN subtypes. Derepression of *Hox13* genes alone does not appear to account for all of the specification defects in *Ring1* mutants, as *Hox10* genes were expressed normally in lumbar segments. Hoxc9 can also repress *Hox4-Hox8* genes in MNs through direct binding (Jung et al., 2010), and likely contributes to the *Ring1* mutant phenotype. Derepression of multiple caudal *Hox* paralogs therefore appears to have a cumulative effect on repressing brachial and thoracic *Hox* genes. In conjunction with studies in *Drosophila*, these finding indicate a deeply-conserved mechanism of rostrocaudal patterning, in which PRCs establish and maintain rostral *Hox* boundaries, while Hox cross-repressive interactions define caudal boundaries.

### PRC2-Independent Functions of PRC1 in Maintaining Neuronal Subtype Fate

Studies of neuronal and non-neuronal development provide compelling evidence that PRC2 activity is required to restrict gene expression in early development (Ezhkova et al., 2009; Gentile et al., 2019; Snitow et al., 2015; Yaghmaeian Salmani et al., 2018). An unexpected finding from our studies is that MN subtype differentiation and *Hox* boundaries are maintained after removal of PRC2 activity. Mutation of *Ezh* genes or *Eed* depletes H3K27me3 from progenitors and postmitotic neurons, without appreciably affecting MN generation, *Hox* expression, or downstream Hox effectors. These observations indicate that neural progenitors can acquire independence from continuous PRC2 activity to restrict gene expression, whereas there is an absolute requirement for PRC1 function to maintain appropriate transcription factor expression and chromatin organization.

How does PRC1 restrict expression of fate determinants in the absence of PRC2 activity? One possibility is that residual H3K27me3, below the threshold of histological detection, is transmitted through cell division and is sufficient to recruit canonical PRC1 and maintain target gene repression. As cells divide, newly synthesized histones are presumably devoid of H3K27me3 in PRC2 mutants, leading to replication-coupled dilution of H3K27me3. During neural differentiation, the rate of cell division decreases (Kicheva et al., 2014; Wilcock et al., 2007), potentially limiting H3K27me3 reduction, and enabling PRC1 to bind at target loci, even in the absence of *de novo* H3K27 methylation. Although we observed no MN phenotypes in *Ezh* or *Eed* mutants, later-born oligodendrocytes that derive from the same precursors as MNs have been shown to depend on PRC2 function for their differentiation (Wang et al., 2020). This more pronounced effect on glial development likely reflects further dilution of H3K27me3 through cell division in PRC2 mutants.

Stabilization of PRC1 at repressed loci could also maintain target repression independently of *de novo* H3K27 methylation. Repression by canonical PRC1 has been shown to depend on the formation of phase-separated subnuclear structures (Polycomb bodies) assembled through polymerization of Polyhomeotic-like (Phc) and/or Cbx proteins (Isono et al., 2013; Plys et al., 2019; Tsuboi et al., 2018). Mutation of *Phc2* in mice leads to ectopic expression of *Hox* genes, with *Hox13* genes among the most robustly derepressed targets (Isono et al., 2013). One possibility is that PcG-mediated repression may not depend on anchoring of PRC1 through H3K27me3, but is maintained through Phc-mediated chromatin compaction at specific loci.

We suggest that in the early phases of development PRC2 activity defines the sites of maintained H3K27 tri-methylation, initiates PRC1 recruitment, and restricts transcription factor expression in MNs. These activities are likely deployed prior to neural progenitor specification, as recent studies have demonstrated that the establishment of rostrocaudal positional identities can be specified before neural induction (Metzis et al., 2018). Patterning morphogens, such as RA and FGF, may initially act on stem cells to establish the pattern of H3K27me3 at *Hox* loci prior to neurogenesis. This early rostrocaudal patterning step may ultimately serve to coordinate *Hox* expression in the neural tube and surrounding mesodermally-derived tissues. The subsequent switch to reliance on PRC1 function could reflect a general mechanism of gene regulation in neuronal subtypes that become terminally differentiated during development.

## Materials and Methods

### Mouse Genetics

*Ezh1 flox* (Hidalgo et al., 2012), *Ezh2 flox* (Su et al., 2003), *Olig2::Cre* (Dessaud et al., 2007), *Hb9::GFP* (Arber et al., 1999) mice have been previously described. *Ring1A−/−; Ring1B flox* (Cales et al., 2008; del Mar Lorente et al., 2000) and *Rybp flox; Yaf2−/−* (Hisada et al., 2012; Rose et al., 2016) mouse lines were generated by microinjection of the mouse ESCs expressing these constructs into the blastocyst followed by implantation of these eggs into pseudopregnant foster female mice. Generation of *Ezh^MNΔ^* mice was performed by crossing *Ezh1 flox, Ezh2 flox* and *Olig2::Cre*. *Ezh^MNΔ^* mice are viable at birth but do not survive beyond P20. Generation of *Ring1^MNΔ^* mice was performed by crossing *Ring1A−/−; Ring1B flox* and *Olig2::Cre*. *Ring1^MNΔ^* mice perish at birth. Generation of *Ring1^MNΔ^*;*Hb9::GFP* mice was performed by crossing *Ring1A−/−; Ring1B flox*, *Olig2::Cre* mice to *Hb9::GFP* mice. Animal work was approved by the Institutional Animal Care and use Committee of the NYU School of Medicine in accordance to NIH guidelines.

### Slide Immunohistochemistry

Embryos were fixed in 4% PFA for 1.5-2 hours at 4°C, washed 5-6 times in cold PBS for 15-30 minutes each wash, and incubated overnight in 30% sucrose. Tissue was embedded in OCT, frozen in dry ice, and sectioned at 16 μm on a cryostat. For antibody staining of sections, slides of cryosections were placed in PBS for 5 minutes to remove OCT. Sections were then transferred to humidified trays and blocked for 20-30 minutes in 0.75 ml/slide of PBT (PBS, 0.1% Triton) containing 1% Bovine serum albumin (BSA). The blocking solution was replaced with primary staining solution containing antibodies diluted in PBT with 0.1% BSA. Primary antibody staining was performed overnight at 4°C. Slides were then washed three times for 5 minutes each in PBT. Fluorophore-conjugated secondary antibodies were diluted 1:500-1:1000 in PBT and filtered through a 0.2 μm syringe filter. Secondary antibody solution was added to slides (0.75 ml/slide) and incubated at room temperature for 1 hour. Slides were washed three times in PBT, followed by a final wash in PBS. Coverslips were placed on slides using 110 μl of Vectashield (Vector Laboratories).

Antibodies against Hox proteins and MN subtypes were generated as described (Dasen et al., 2008; Dasen et al., 2005). Additional antibodies were used as follows: goat anti-Ring1B (Abcam, 1:2000), rabbit anti-Ring1B (Abcam, 1:5000), rabbit anti-Rybp (Abcam1:2000), rabbit anti-Yaf2 (Abcam, 1:2000), rabbit anti-Cbx2 (Bethyl, 1:5000), rabbit anti-H3K27me3 (Cell Signaling, 1:2000).

### *In situ* mRNA Hybridization

Probe templates were generated by RT-PCR and incorporated a T7 promoter sequence in the antisense strand. Total RNA was first extracted from eviscerated E12.5 embryos using TRIzol (Invitrogen). Genes of interest were amplified with the One Taq One-Step RT-PCR kit (NEB) using 1 μg of RNA. After amplification by RT-PCR, a second PCR was performed to incorporate a T7 promoter sequence. Antisense riboprobes were generated using the Digoxigenin-dUTP (SP6/T7) labeling kit (Sigma-Aldrich). For *in situ* hybridization, sections were first dried for 10-15 minutes at room temperature, placed in 4% PFA, and fixed for 10 minutes at room temperature. Slides were then washed three times for 3 minutes each in PBS, and then placed in Proteinase K solution (1 μg/ml) for 5 minutes at room temperature. After an additional PFA fixation and washing step, slides were treated in triethanolamine for 10 minutes, to block positive charges in tissue. Slides were then washed three times in PBS and blocked for 2-3 hours in hybridization solution (50% formamide, 5X SSC, 5X Denhardt’s solution, 0.2 mg/ml yeast RNA, 0.1 mg/ml salmon sperm DNA). Prehybridization solution was removed, and replaced with 100 μl of hybridization solution containing 100 ng of DIG-labeled antisense probe. Slides were then incubated overnight (12-16 hours) at 72°C. After hybridization, slides were transferred to a container with 400 ml of 5X SSC and incubated at 72°C for 20 minutes. During this step, coverslips were removed using forceps. Slides were then washed in 400 ml of 0.2X SSC for 1 hour at 72°C. Slides were transferred to buffer B1 (0.1 M Tris pH 7.5, 150 mM NaCl) and incubated for 5 minutes at room temperature. Slides were then transferred to staining trays and blocked in 0.75 ml/slide of B1 containing 10% heat inactivated goat serum. The blocking solution was removed and replaced with antibody solution containing 1% heat inactivated goat serum and a 1:5000 dilution of anti-DIG-AP antibody (Roche). Slides were then incubated overnight at 4°C in a humidified chamber. The following day, slides were washed 3 times, 5 minutes each, with 0.75 ml/slide of buffer B1. Slides were then transferred to buffer B3 (0.1 M Tris pH 9.5, 100 mM NaCl, 50 mM MgCl_2_) and incubated for 5 minutes. Slides were then developed in 0.75 ml/slide of B3 solution containing 3.5 μl/ml BCIP and 3.5 μl/ml NBT for 12-48 hours. After color development, slides were washed in ddH_2_0 and coverslipped in Glycergel (Agilent). A more detailed in situ hybridization protocol is available on our lab website (http://www.dasenlab.com).

### Wholemount Immunohistochemistry

For wholemount immunohistochemistry embryos were fixed in PFA for 2 hours, then bleached for 24 hours at 4°C in a 10% H_2_O_2_, 10% DMSO solution prepared in methanol. Embryos were washed three times for 10 minutes each in methanol, followed by five washes for 10 minutes in PBS. Primary antibodies were diluted in staining solution (5% BSA, 20% DMSO in PBS) and specimens were incubated in staining solution on a rotator overnight at room temperature. Samples were then washed three times for 5 minutes each in PBS, followed by four 1 hour washes in PBS. Specimens were then incubated in secondary antibodies diluted in staining solution overnight at room temperature. Samples were then washed three times for 5 minutes each in PBS, followed by four 1 hour washes in PBS, a single 10 min wash in 50% methanol, and three 20 minute washes in 100% methanol. Samples were transferred to glass depression slides and tissue was cleared by incubating samples in BABB solution (1-part benzyl alcohol: 2-parts benzyl benzoate). Confocal images of embryos were obtained from Z-stacks using Zen software (Zeiss). Further details of wholemount staining protocols are available on our lab website: (http://www.dasenlab.com).

### *In ovo* chick electroporation

*In ovo* electroporation were performed on Hamburger Hamilton (HH) stage 13-14 chick embryos and analyzed at HH stage 24-25. Fertilized chicken eggs (Charles River) were incubated in a humidified incubator at 39°C for 40-48 hours until they reached HH13-14. The top of the egg shell was removed and a 1 μg/μl DNA (150-500 ng/μL expression plasmid and pBKS carrier DNA) containing ~0.02% Fast green was injected into the central canal of the neural tube using a sharpened glass capillary tube. Electrodes (Platinum/Iridium (80%/20%), 250 μm diameter, UEPMGBVNXNND, FHC Inc.) were placed on both sides of the neural tube (4 mm separation) and DNA was electroporated using an ECM 830 electroporator (ECM 830, BTX; 25V, 4 pulses, 50 ms duration, 1 second interval). Eggs were sealed with parafilm and incubated for 48 hours prior to fixation. Results shown in figures are representative of at least 3 electroporated embryos from two or more experiments in which electroporation efficiency in MNs was above 60%.

### RNA preparation and Library Preparation

RNA was extracted from FACS purified MNs dissected from E12.5 mouse embryo (10-20,000 cells/segment), using the Arcturus Picopure RNA isolation kit. For on-column DNase treatment, Turbo DNase was used (ambion, AM2238). Each samples used separated bar code for libraries. RNA quality and quantity were measured with an Agilent Picochip using a Bioanalyzer, all samples had quality scores between 9-10 RIN. For library preparation 10ng of total RNA was used to generate cDNA, which was amplified with SMARTer Stranded RNA-Seq kit. 100ng of cDNA were used as input to prepare the libraries (Takara, #634837), and amplified by 10 PCR cycles. Samples were run in four 50-nucleotide paired-end read rapid run flow cell lanes with the Illumina HiSeq 4000 sequencer.

### RNAseq Data Analyses

Sequencing reads were mapped to the assembled reference genome (mm10) using the STAR aligner (v2.5.0c) (Dobin et al., 2013). Alignments were guided by a Gene Transfer Format (GTF) file. The mean read insert sizes and their standard deviations were calculated using Picard tools (v.1.126) (http://broadinstitute.github.io/picard). The read count tables were generated using HTSeq (v0.6.0) (Anders et al., 2015), normalized based on their library size factors using DEseq2(Love et al., 2014), and differential expression analysis was performed. The Read Per Million (RPM) normalized BigWig files were generated using BEDTools (v2.17.0) (Quinlan and Hall, 2010) and bedGraphToBigWig tool (v4). To compare the level of similarity among the samples and their replicates, we used two methods: principal-component analysis and Euclidean distance-based sample clustering. All the downstream statistical analyses and generating plots were performed in R environment (v3.1.1) (https://www.r-project.org/).

### ATACseq

ATACseq was performed following previously described protocols(Buenrostro et al., 2015). DNA was extracted from purified dissected mouse embryonic MNs. Cells were aliquoted and washed twice in ice-cold 1× PBS. Cell pellets were resuspended in 10 mM Tris (pH 7.4), 10 mM NaCl, 3 mM MgCl2, 0.1% NP-40 (v/v), 0.1% tween20, 0.01% Digitonin and 1% BSA, centrifuged at 500 g for 5 min at 4°C. Pellets were resuspended in 12.5 μl of 2× tagmentation DNA buffer, 1.25 μl Tn5 (Nextera DNA Sample Preparation Kit, FC-121–1030) and 11.25 μl of water, and incubated at 37°C for 30 min. The sample was purified using the MinElute PCR Purification Kit (Qiagen, 28004). PCR enrichment of the library was performed with custom-designed primers and 2× NEB Master Mix. A qPCR reaction with 1× SYBR Green (Invitrogen), custom-designed primers and 2× NEB Master Mix (New England Labs, M0541) was performed to determine the optimal number of PCR cycles (one third of the maximum measured fluorescence)(Buenrostro et al., 2013). The libraries were purified using the AMPure XP beads (Beckman Coulter, A63880). High Sensitivity DNA ScreenTape (Agilent, 5067-5584) was used to verify the fragment length distribution of the library. Library quantification was performed using the KAPA Library Amplification kit on a Roche LightCycler 480. The libraries were sequenced on an Illumina NovaSeq (100 cycles, paired-end).

### ATACseq data analysis

All of the reads from the Sequencing experiment were mapped to the reference genome (mm10) using the Bowtie2 (v2.2.4)(Langmead and Salzberg, 2012) and duplicate reads were removed using Picard tools (v.1.126) (http://broadinstitute.github.io/picard/). Low quality mapped reads (MQ<20) were removed from the analysis. The read per million (RPM) normalized BigWig files were generated using BEDTools (v.2.17.0) (Quinlan and Hall, 2010) and the bedGraphToBigWig tool (v.4). Peak calling was performed using MACS (v1.4.2)(Zhang et al., 2008) and peak count tables were created using BEDTools. Differential peak analysis was performed using DESeq2 (Love et al., 2014). ChIPseeker (v1.8.0)(Yu et al., 2015) R package was used for peak annotations and motif discovery was performed using HOMER (v4.10)(Heinz et al., 2010). ngs.plot (v2.47) and ChIPseeker were used for TSS site visualizations and quality controls. KEGG pathway analysis and Gene Ontology (GO) analysis was performed using the clusterProfiler R package (v3.0.0)(Yu et al., 2012). To compare the level of similarity among the samples and their replicates, we used two methods: principal-component analysis and Euclidean distance-based sample clustering. The downstream statistical analyses and generating plots were performed in R environment (v3.1.1) (https://www.r-project.org/).

### Statistics

Samples sizes were determined based on previous experience and the number of animals and definitions of N are indicated in the main text and figure legends. In figures where a single representative image is shown, results are representative of at least two independent experiments, unless otherwise noted. No power analysis was employed, but sample sizes are comparable to those typically used in the field. Data collection and analysis were not blind. Graphs of quantitative data are plotted as means with standard error of mean (SEM) as error bars, using Prism 8 (Graphpad) software. Unless noted otherwise, significance was determined using unpaired t-test in Prism 8 software, or using adjusted p-values. Exact p-values are indicated, where appropriate, in the main text, figures, and figure legends.

## Data Availability

RNAseq and ATACseq data are available through GEO (GSE175503).

## Acknowledgements

We thank Kristen D’Elia, Sara Fenstermacher, Jessica Treisman, and Ed Ziff for discussion and comments on the manuscript, and Rachel Kim and Orly Wapinski for assistance. We thank Isabel Hidalgo and Susana Gonzalez for providing *Ezh1flox* mice, Alexander Tarakhovsky for *Ezh2 flox* mice, Stefan Thor for *Eed* mutant embryos, Haruhiko Koseki for *Ring1A−/−;Ring1B flox* ES cells, and Robert Klose for *Rybp flox; Yaf2−/−* ES cells. We also thank Genome Technology Center and Cytometry and Cell Sorting Laboratory at NYU Langone. This work was supported by NIH NINDS grants T32 GM007238, F31 NS087772 to AS, R01NS100897 to EOM, R35 NS116858, R01 NS062822 and R01 NS097550 to JD.

## Author Contributions

A.S. and J.S.D. designed experiments, interpreted data, and wrote the paper. S.P. performed in situ hybridizations. A.M. and A.S. performed chick neural tube experiments. M.B. and E.O.M designed and performed mouse ESC experiments. A.S. performed all mouse genetic analyses and performed RNAseq and ATACseq. A. K.-J. analyzed ATACseq and RNAseq data.

---

**Supplementary Figure 1.**
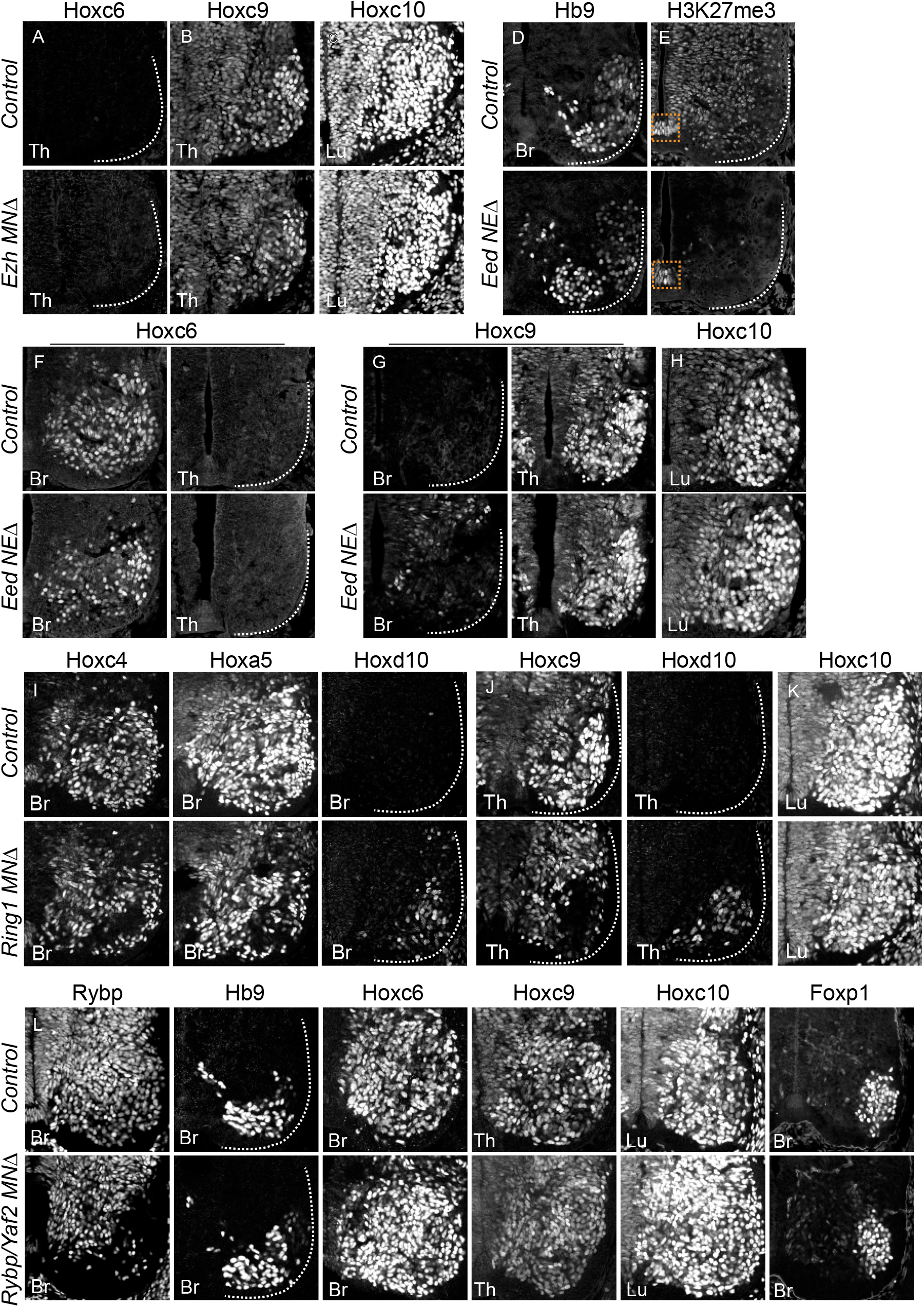
Effects of PRC mutations on MN differentiation. (A-C) Expression of Hoxc9 in thoracic segments and Hoxc10 in lumbar segments is normal in E11.5 *Ezh^MNΔ^* mice (*Eed flox/flox, Sox1::Cre/*^+^), and no ectopic thoracic Hoxc6 expression is detected. (D,E) In *Eed^NEΔ^* mice (H3K27me3 is depleted throughout the spinal cord at E12.5, while Hb9 expression is unaffected. Some H3K27me3 is detected in the floor plate of *Eed^NEΔ^* mice (boxed region). (F-H) Brachial Hoxc6, thoracic Hoxc9, and lumbar Hoxc10 are detected normally in *Eed^NEΔ^* mice. (I) Brachial Hoxc4 and Hoxa5 are lost from MNs in *Ring1^MNΔ^* mice, and Hoxd10 is ectopically expressed in brachial MNs. (J) Hoxc9 expression is reduced in thoracic segments of *Ring1^MNΔ^* mice, and Hoxd10 is expressed. (K) Expression of Hoxc10 is retained at lumber levels of *Ring1^MNΔ^* mice. (L) Analysis of E12.5 *Rybp/Yaf2MNΔ* mice (*Rybp ^flox/flox^; Yaf2^−/−^; Olig2::Cre*/^+^). Rybp expression is selectively depleted from MNs. Hb9 and indicated Hox proteins are expressed normally.

**Supplementary Figure 2.**
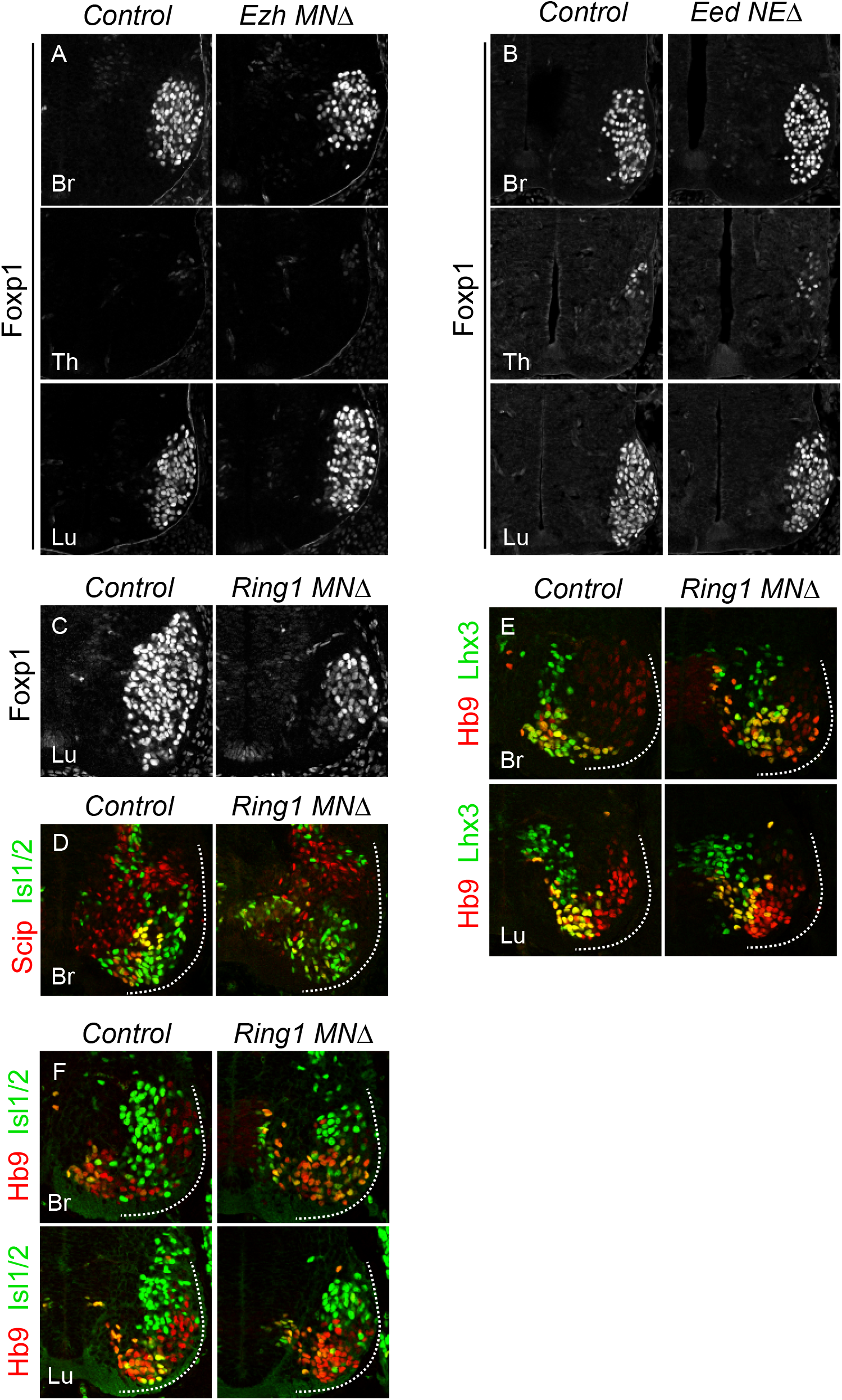
Analyses of MN subtypes in PRC mutant mice. (A,B) Foxp1 expression is unaffected at brachial, thoracic, and lumbar levels in *Ezh^MNΔ^* and *Eed^NEΔ^* mice. (C) Reduced Foxp1 expression in lumbar segment of *Ring1^MNΔ^* mice. (D) Reduced expression of phrenic motor column (PMC) markers (Scip^+^, Isl1/2^+^) in rostral brachial segments of *Ring1^MNΔ^* mice. (E) Expression of MMC neuron (Hb9, Lhx3) markers in brachial and lumbar segments of *Ring1^MNΔ^* mice. (F) Expression of HMC markers (Hb9, Isl1/2) in brachial and lumbar segments of *Ring1^MNΔ^* mice.

**Supplementary Figure 3.**
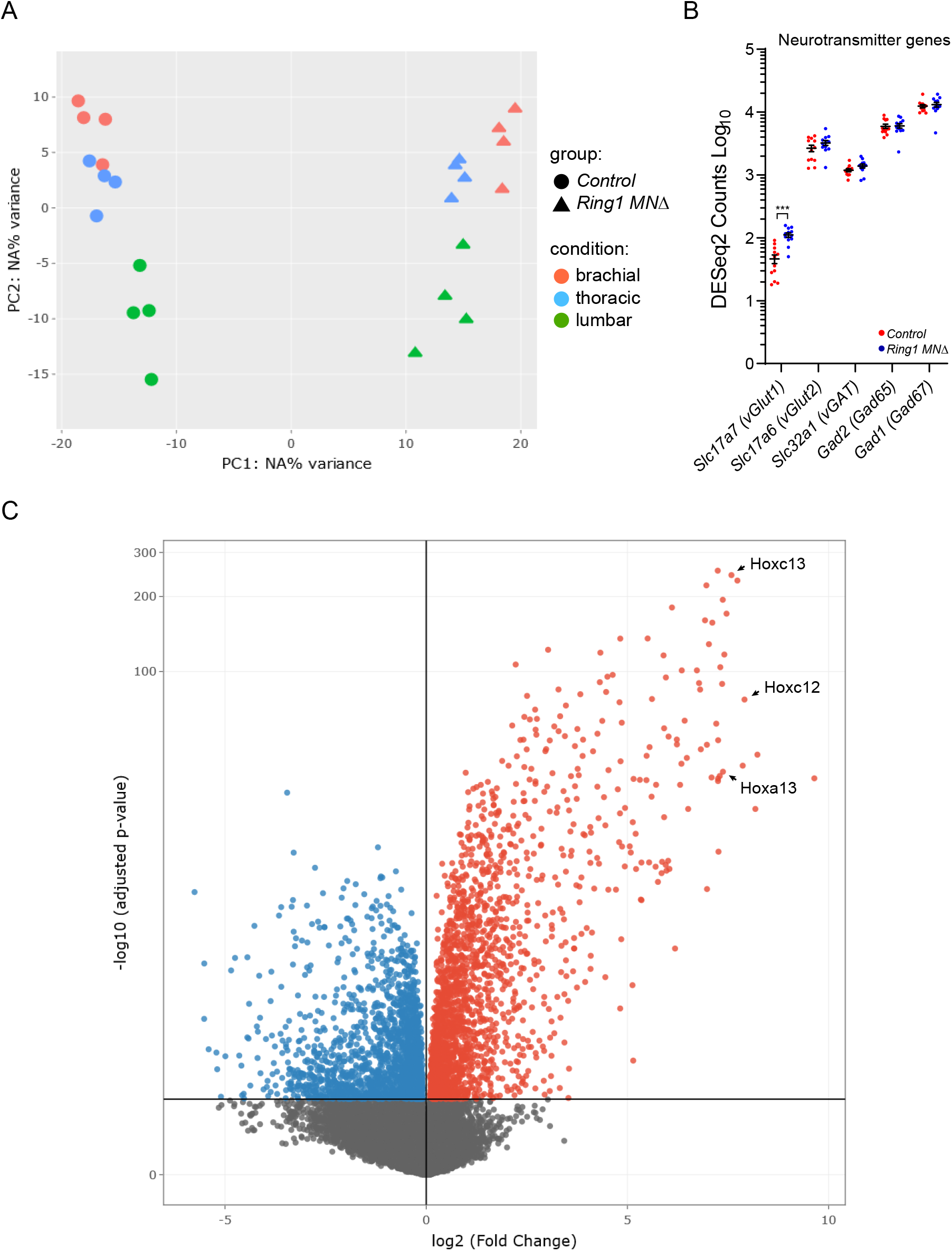
RNAseq analyses of Ring1 mutant mice. (A) PCA plot of RNAseq samples. (B) Plot of neurotransmitter encoding genes in control and *Ring1^MNΔ^* mice. Commonly used gene names are shown in parenthesis. Expression of *Slc17a7* is increased in *Ring1^MNΔ^* mice, although the absolute DESeq2 counts were still relatively low (46.3 +/− 7.2 in controls versus 111.1 +/− 9 in *Ring1^MNΔ^* mice, p=0.000012). Elevated expression of *Gad1* in controls and mutants may be due to presence of spinal interneurons in sorted samples. (C) Volcano plot of upregulated (red) and downregulated (blue) genes. Three caudal *Hox* genes are indicated.

**Supplementary Figure 4.**
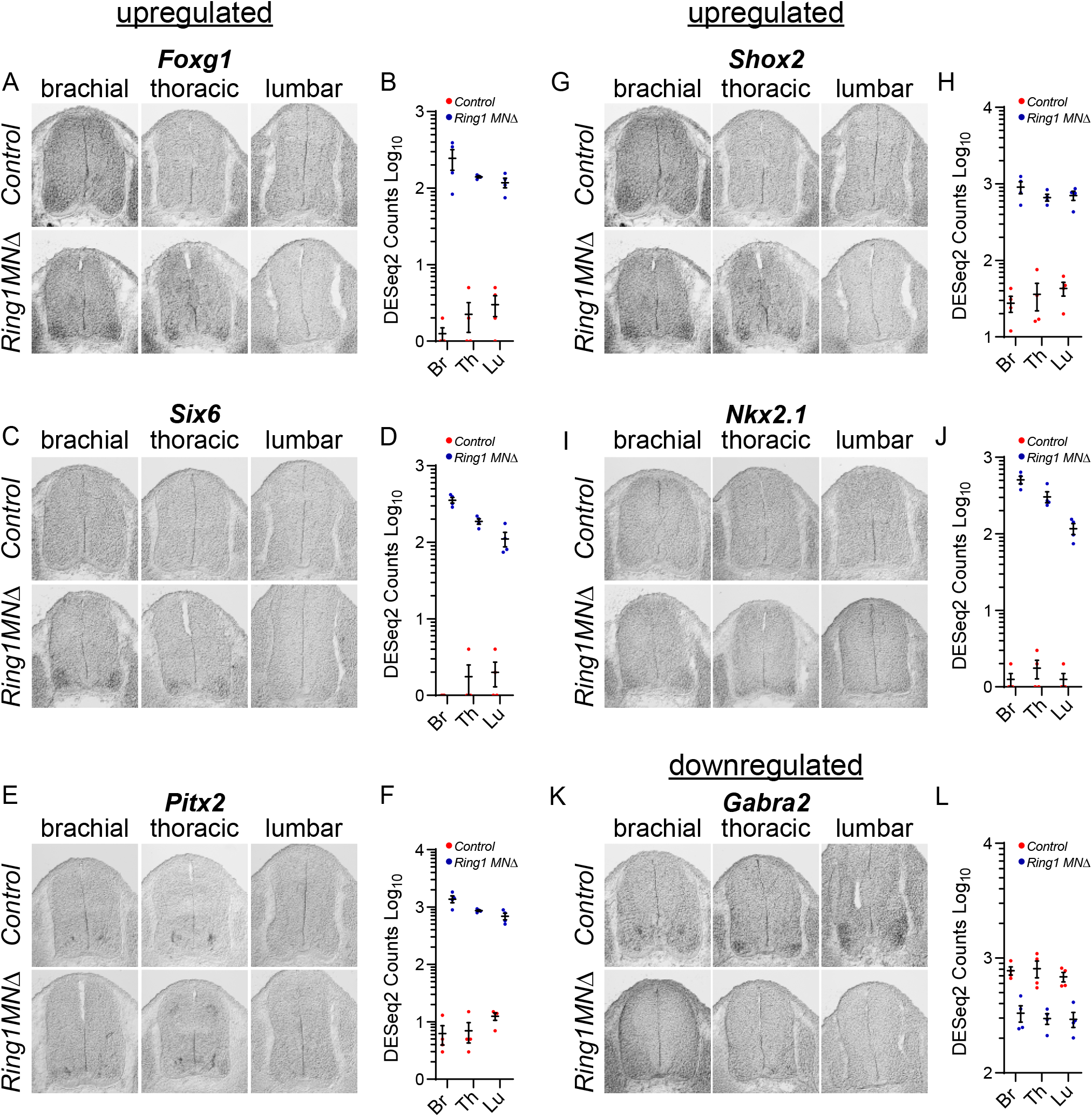
Validation of downregulated genes from *Ring1^MNΔ^* RNAseq. (A,C,E,G,I) *In situ* hybridization of selected upregulated genes from the *Ring1^MNΔ^* RNAseq. Images show sections at brachial, thoracic, and lumbar levels from E12.5 control and *Ring1^MNΔ^* mice. (B,D,F,H,J) Graphs of DESeq2 counts for each upregulated gene in each segments. (K) Analysis of the downregulated gene *Gabra2* by *in situ* hybridization. (L) Graph of DESeq2 counts for *Gabra2* in each segment.

**Supplementary Figure 5.**
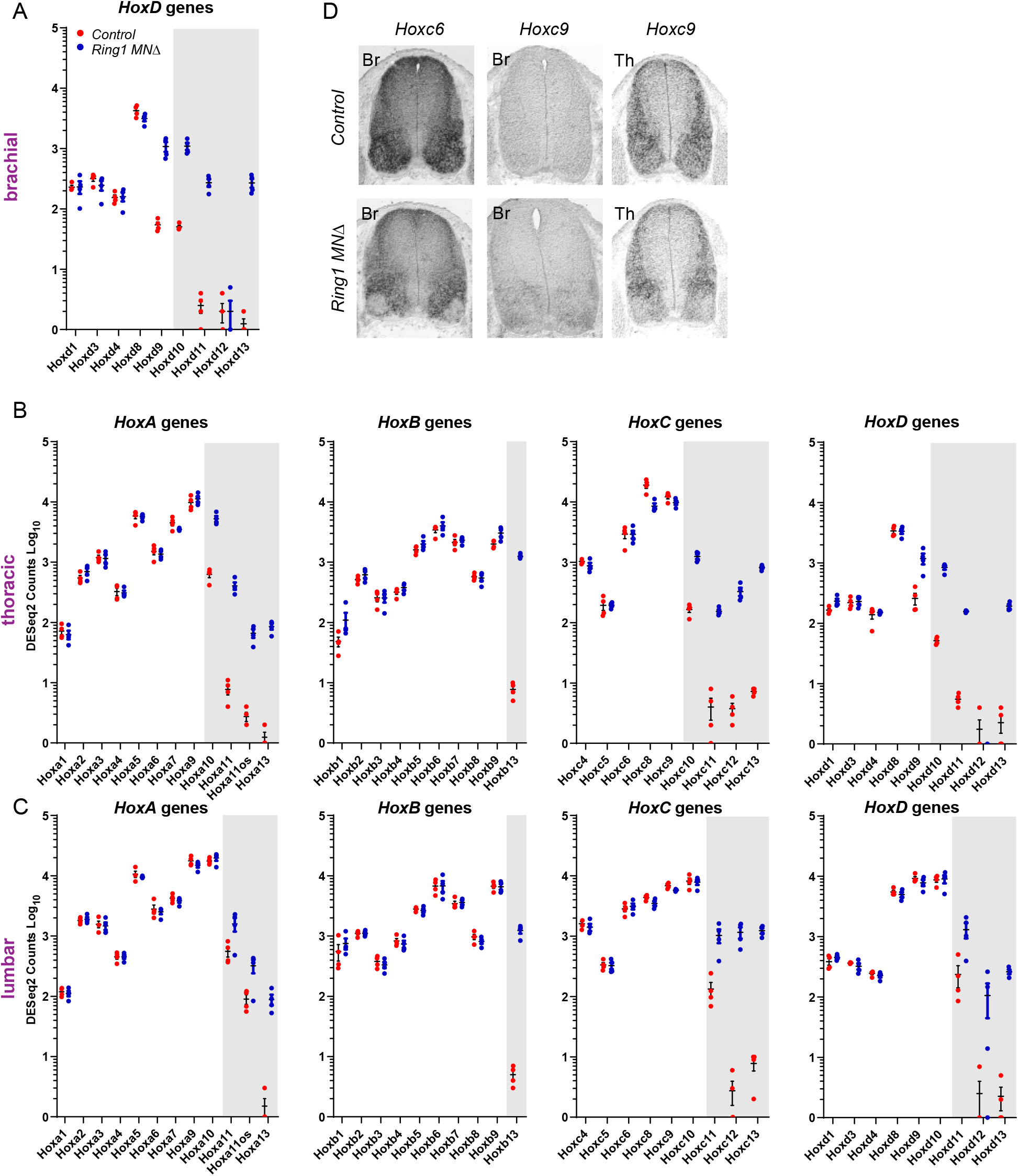
Derepression of caudal *Hox* genes in *Ring1^MNΔ^* mice. (A-C) DESeq2 counts of *HoxA*, *HoxB*, *HoxC*, and *HoxD* gene clusters in brachial (A), thoracic (B), and lumbar (C) segments. Gray-shaded regions highlight *Hox* genes that are derepressed in MNs of *Ring1^MNΔ^* mice. In lumbar segments, expression of *Hox10* genes is not reduced in *Ring1* mutants, although *Hox13* genes are derepressed. (D) *In situ* hybridization of *Hoxc6* and *Hoxc9* in *Ring1^MNΔ^* mice at brachial (Br) and thoracic (Th) levels at E12.5. In brachial segments, expression of *Hoxc6* is lost from MNs, and *Hoxc9* is expressed, similar to the analyses of Hox protein expression. In thoracic segments, *Hoxc9* expression is reduced in MNs.

**Supplementary Figure 7.**
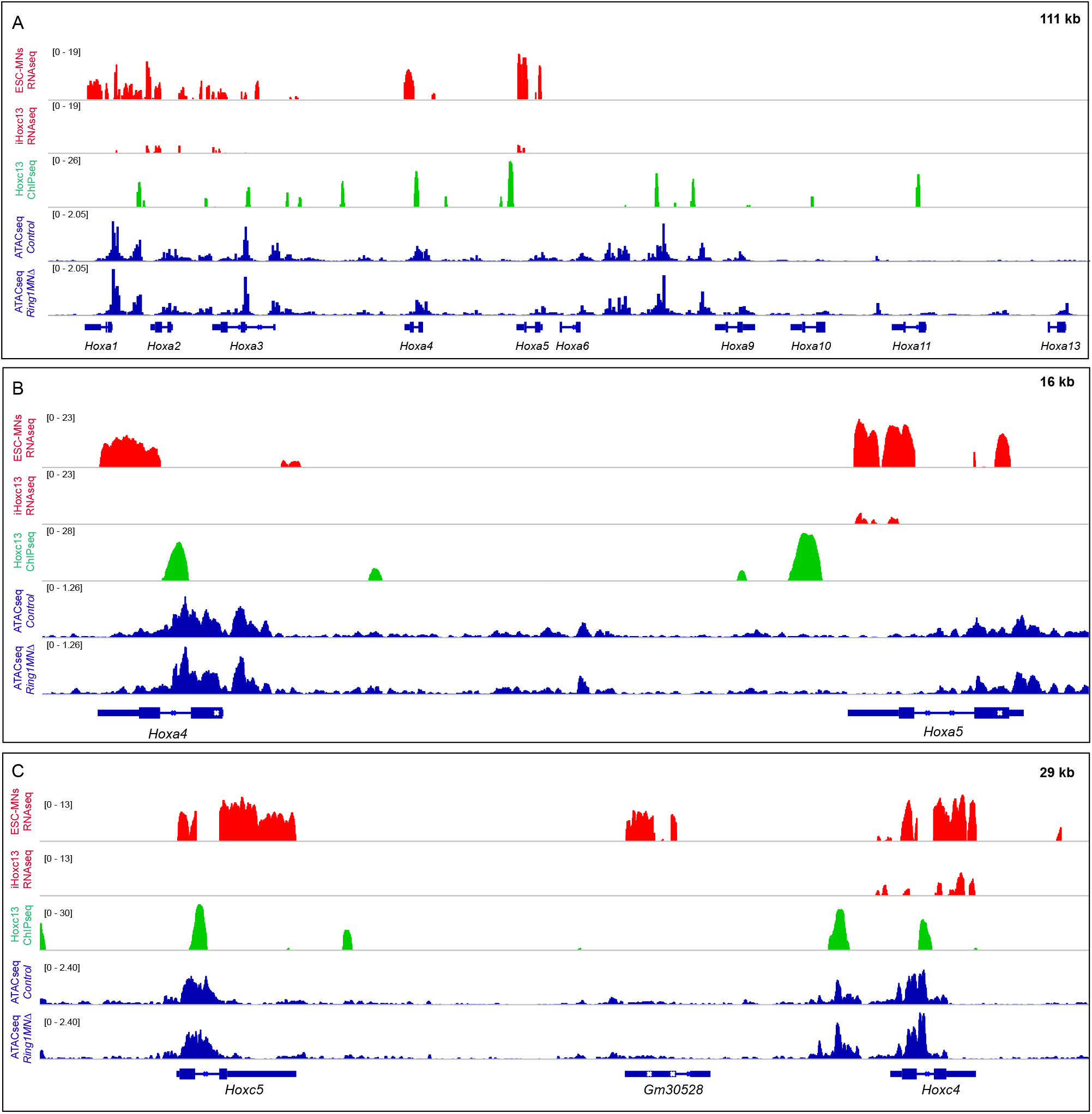
Comparison of Hoxc13 binding and accessibility in control and *Ring1^MNΔ^* MNs. (A) Effects of *Hoxc13* expression in ESC-MNs at the *HoxA* locus. Top panels show IGV browser views (log scale) of RNAseq (red) and ChIPseq tracks (green). Induced Hoxc13 represses expression of *Hoxa1*-*Hoxa5* genes and binds near multiple *HoxA* genes. Bottom panels show ATACseq in brachial control and *Ring1* mutant MNs at the *HoxA* cluster (blue). Bottom panels show ATACseq in brachial control and *Ring1* mutant MNs at the *HoxA* cluster. (B,C) Magnified views of *Hoxc13* binding, RNA expression, and ATACseq results at *Hox4-Hox5* genes in the *HoxA* and *HoxC* clusters.

